# The nuclear Argonaute HRDE-1 broadly interacts with euchromatin via an RGG domain to enable transgenerational epigenetic silencing

**DOI:** 10.64898/2026.10.15.759339

**Authors:** Julie Z. Ni, Natasha Mason, Junjie Liu, Meghana Thimmaraju, Brina Trivedi, Joyce Naguib, Sam G. Gu

## Abstract

Small RNA(sRNA)-guided transcriptional repression, a process termed as nuclear RNAi, safeguards genome integrity and germline development. Central to this pathway are the highly conserved Argonaute (AGO) proteins, which are guided by sRNAs to mediate transcriptional silencing and heterochromatin formation at the target locus. The mechanism of how nuclear AGO proteins engage chromatin remains poorly understood. By studying *C. elegans* nuclear AGO HRDE-1, we found that an intrinsically disordered arginine-glycine-glycine (RGG) domain in HRDE-1 is essential for siRNA-guided transgenerational epigenetic silencing and germline immortality. The function of RGG occurs in the downstream of siRNA loading and nuclear localization. Mutant HRDE-1 that lacks the RGG domain is defective in binding to the chromatin of the siRNA targets. Strikingly, ChIP-seq analysis reveals that HRDE-1 interacts with hundreds of highly transcribed euchromatic protein-coding genes in the germ cells, which is unexpected for an epigenetic silencing protein. The HRDE-1-euchromatin interaction is not mediated by siRNAs but requires the RGG domain. The interaction appears to be necessary but not sufficient to cause gene silencing. We propose that the initial siRNA-independent, RGG-dependent HRDE-1-chromatin interaction enables HRDE-1 to enter the transcribing chromatin compartment, a likely prerequisite for the subsequent siRNA-pre-mRNA interaction. Given the prevalence of the RGG domain in eukaryotic proteins, we constructed chimera HRDE-1 proteins and found that RGGs from other proteins, including human hnRNP U, can substitute for the HRDE-1 RGG to restore chromatin binding and transgenerational silencing. Our findings suggest an evolutionarily conserved role for RGG-rich intrinsically disordered regions in coupling AGOs and other RNA-binding proteins to active chromatin as a mechanism to target nascent pre-mRNA.

## Introduction

RNA interference (RNAi) was initially discovered as a gene silencing activity triggered by exogenous homologous dsRNA ^1,2^ or transgene ^3–5^. A central component of RNAi is the Argonaute (AGO) protein, which exists in all three domains of life. The canonical eukaryotic AGO proteins target RNA molecules to mediate gene silencing (reviewed in ^6–8^). Structurally, AGOs share a bilobed N-PAZ-MID-PIWI architecture that binds a 20-30-nt long guide RNA molecule and can accommodate the base-pairing interactions between the guide RNA and target RNA (or DNA in some cases) (reviewed in ^9^). In the classical RNAi pathway, AGO degrades the target transcript by the “slicer” activity of the PIWI domain ^10^. For the microRNAs pathway, AGO can either degrade the target RNA or repress protein translation through associated proteins (reviewed in ^11^). In addition to these post-transcriptional gene silencing mechanisms, AGO can induce a long-term gene silencing at the chromatin level by heterochromatin formation and transcriptional repression, which is referred to as RNA-induced transcriptional silencing (RITS) or nuclear RNAi (reviewed in ^12–16^). The main function of nuclear RNAi is to silence transposons and other types of DNA repeats in eukaryotic genomes, therefore essential for the genome integrity. Since its discovery, nuclear RNAi has been recognized as a key model system to explore RNA-mediated epigenetic mechanisms that connect genomic, developmental, and environmental signals with gene expression.

The following is a simplified model of how nuclear RNAi works. Despite of a repressive state, a nuclear RNAi targeted region is transcribed at a low level to produce either small interfering RNA (siRNAs) ^17^ or piRNAs ^18^, which are loaded into the AGO or PIWI clades of the AGO proteins, respectively, depending on the species. These AGO or PIWI proteins form the RITS complex with other proteins including chromatin modifying enzymes and RNA processing enzymes. The siRNAs or piRNAs then guide the RITS to the target nascent transcripts to maintain the repressive chromatin state by depositing DNA or/and histone modifications that lead to heterochromatin formation (reviewed in ^12,14,19^).Among different species that have nuclear RNAi, *C. elegans* stands out as a uniquely powerful model organism to study this pathway because of several distinctive advantages. First, unlike many other organisms in which AGO functions in both cytoplasm and nucleus, *C. elegans* possesses dedicated nuclear AGO proteins. For example, *Drosophila* AGO2 ^20–22^ and human AGO1 and AGO2 ^23–25^ function in both the cytoplasm and nucleus, making it less straightforward to dissect their nuclear specific function. In contrast, the *C. elegans* nuclear AGO proteins, HRDE-1 ^26–28^ and NRDE-3 ^29^, are dedicated to nuclear RNAi. NRDE-3 primarily functions in the somatic cells ^29^, although a low germline expression has been observed for NRDE-3 ^30^, whereas HRDE-1, which is the focus of this study, is a germline-specific nuclear AGO ^30^. Second, *C. elegans* offers exceptional genetic tractability. Mutant strains that are defective in germline nuclear RNAi are fertile at the permissive temperature ^26^ and are readily available for various functional and molecular studies. Third, germline nuclear RNAi can be readily induced at endogenous loci using exogenous dsRNA or piRNA triggers, resulting in robust transcriptional repression and deposition of heterochromatin marks (H3K23me3, H3K27me3, and H3K9me3) ^26–28,31–35^. Remarkably, both the transcriptional repression and heterochromatin marks can persist for multiple generations after the initial dsRNA or piRNA trigger has been removed ^26,31,34,36^, providing an experimentally tractable system for investigating transgenerational epigenetic inheritance (TEI). Finally, germline nuclear RNAi is linked to germline immortality. Loss-of-function mutant animals in this pathway exhibit a temperature-sensitive fertility defect. Although fertile at permissive temperature, they become progressively sterile after several generations at 23-25°C, which is well tolerated by the wild-type animal. This phenotype, known as the mortal germline or Mrt phenotype ^26,37^, is likely caused by cumulative epigenetic defects ^38,39^. Since the fertility defect is transgenerational cumulative and temperature sensitive, germline nuclear RNAi in *C. elegans* has been used as a tractable system to study the impact of ancestral stress to the offspring epigenome and reproduction.

Despite recent progress in nuclear RNAi, the molecular mechanism that leads to chromatin targeting and silencing are largely uncharacterized. In the simplest model, a nuclear AGO such as HRDE-1 is recruited to a transcribing chromatin through base-pairing interactions between siRNA and chromatin-bound nascent transcript. Yet, some of the fundamental and pressing questions remain unanswered: How does a siRNA-loaded nuclear AGO recruited to chromatin-bound pre-mRNA rather than the mature mRNA? Does nuclear AGOs reach chromatin by random diffusion, or is its recruitment facilitated by other unknown mechanism? It has been proposed that actively transcribing chromatin is organized into biomolecular condensates, forming a compartment physically distinct from the surrounding nucleoplasm (reviewed in ^40^). *In vitro* studies have demonstrated that biomolecular condensate can be formed by multivalent weak interactions between RNA and intrinsically disordered regions of transcription ^41^ and mRNA splicing machineries ^42^, as well as nucleosomes ^43^. Such compartment potentially poses a barrier for AGO to bind a chromatin-bound nascent transcript. Therefore, additional components other than siRNA may be required to facilitate nuclear AGO to enter the transcribing chromatin compartment.

Growing evidence indicates that intrinsically disordered regions and low-complexity motifs, including arginine-glycine-glycine/arginine-glycine (RGG/RG) repeats, can mediate RNA binding, chromatin association, and liquid-liquid phase separation in RNA-binding proteins (reviewed in ^44,45^). The RGG domain is defined by at least two RGG/RG repeats separated by only a few other amino acid residues. More than a thousand human proteins, many of which are RNA-binding proteins, contain the RGG domain. The guanidino group of arginine can bind to an RNA or protein partner via several mechanisms: electrostatic interactions, hydrogen bonding, hydrophobic interactions, and a pi (π) electron orbital that can favor π-π stacking and π-cation interactions. The glycine in RGG/RG can provide the conformational flexibility as well as π stacking and hydrogen bonding for RNA or protein interactions. The RGG domain appears to be able to bind a wide range of RNA sequences, with the interaction to be highly dynamic, multivalent, and low affinity. The importance of the RGG domain has been demonstrated in disease genes such as hnRNP U ^46,47^, FUS ^48^, and fibrillarin ^49^. Studies in *C. elegans* have shown that the RGG domain contributes to the assembly of germline condensates like P granules (reviewed in ^50^) and nucleolar subdomains ^51^, linking RNA metabolism to nuclear organization.

This study demonstrates that a previously uncharacterized RGG domain in HRDE-1 act as a chromatin-engagement module that enables broad access to actively transcribed euchromatin, functioning alongside, but independently of, small-RNA-guided targeting activity.

## Results

### HRDE-1 binds siRNA-targeted heterochromatin

To define how HRDE-1 associates with chromatin, we performed anti-FLAG ChIP-seq using a strain in which endogenous HRDE-1 is tagged with GFP::3×FLAG at the N-terminus. This strain was previously created and confirmed for its wild-type HRDE-1 function ^30^. Anti-FLAG ChIP from two biological replicates of GFP::3×FLAG::HRDE-1 yielded highly similar genome-wide profiles (Fig. 1A). To identify HRDE-1-enriched regions, we performed differential ChIP-seq analysis (DEseq2 ^52^) in 1kb genomic tiling windows, using two different controls: the ChIP input of the GFP::3×FLAG::HRDE-1 strain or anti-FLAG ChIP-seq of an untagged wild-type N2 strain. Using a ≥1.5-fold enrichment threshold and FDR ≤ 0.05, we identified 4512 kb and 2568 kb of genomic regions enriched over input or over anti-FLAG ChIP from untagged N2, respectively, with 1792 kb (1.8% of the genome) called in both comparisons (hypergeometric p-value for the overlap = 0; Fig. 1B and C). These analyses indicate that HRDE-1 interacts with germline chromatin at specific genomic loci. For simplicity, we will use the phrase HRDE-1 ChIP-seq for the rest of the paper to refer to the anti-FLAG ChIP-seq experiment using the GFP::3×FLAG::HRDE-1.

**Figure 1.**
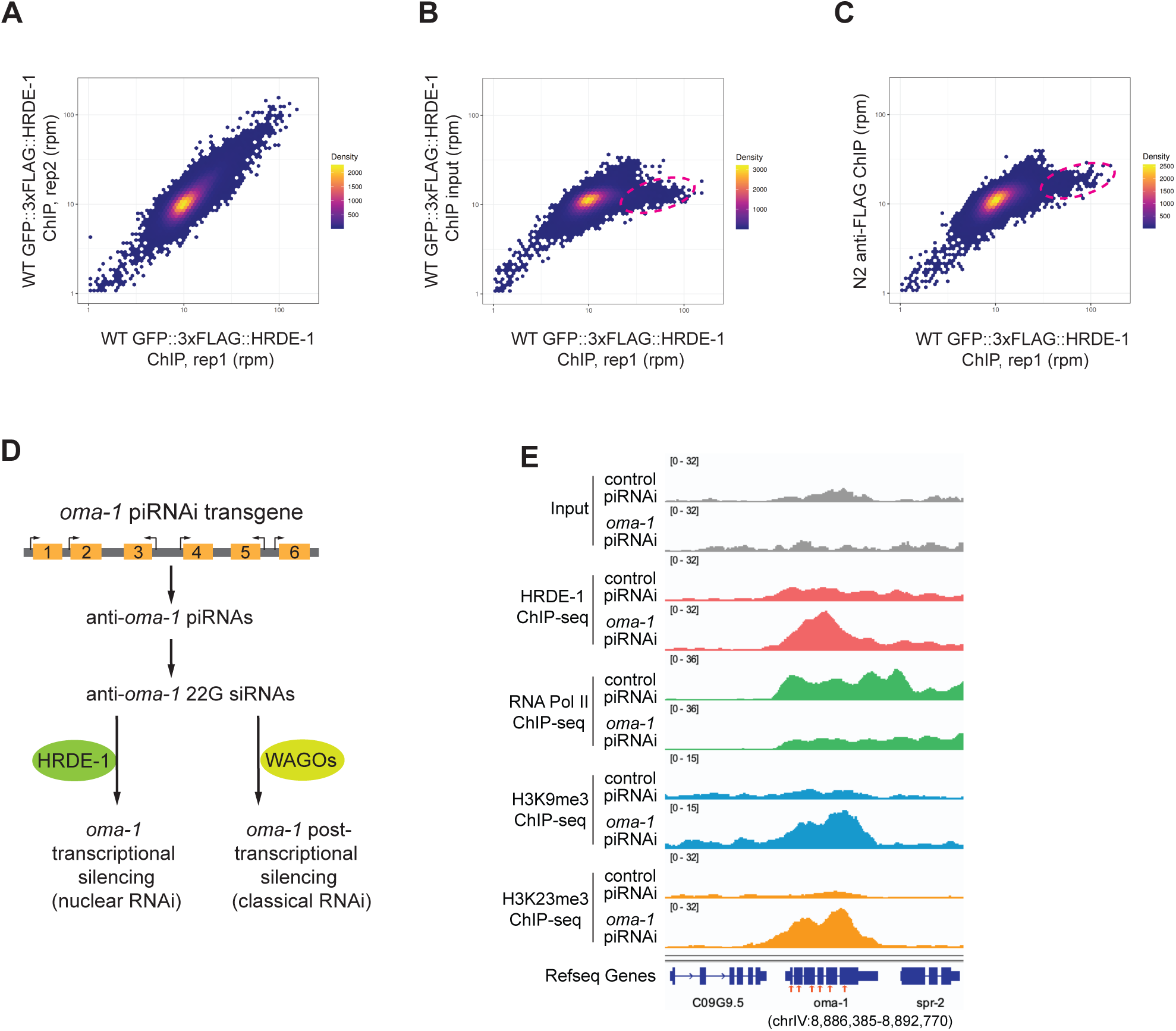
HRDE-1 ChIP-seq reveals specific HRDE-1-chromatin interactions. (A-C) Density scatter plots comparing (A) two biological replicates of GFP::3×FLAG::HRDE-1 ChIP-seq, (B) GFP::3×FLAG::HRDE-1 ChIP-seq and input, and (C) GFP::3×FLAG::HRDE-1 ChIP-seq and control ChIP (anti-FLAG of the N2 sample). RPM values for all 1kb windows in the genome were calculated and used for the plots. (D) Schematic diagram of anti-*oma-1* piRNAi used to induce nuclear RNAi at the *oma-1* locus. (E) ChIP-seq coverage plots at the *oma-1* locus.

We next asked whether HRDE-1 is recruited to a germline expressed gene targeted for nuclear RNAi. The germline-expressed gene *oma-1* is a well-established target to study gene silencing induced by exogenous dsRNA ^53^ or piRNA ^31^. dsRNA or piRNAi against *oma-1* induces HRDE-1-dependent transcriptional repression and heterochromatin formation at this locus ^31,54^. We expressed six artificial anti-*oma-1* piRNAs from a piRNAi transgene and confirmed nuclear RNAi by increased H3K9me3 and H3K23me3 and reduced RNA polymerase II occupancy at *oma-1* relative to a control piRNAi strain (Fig. 1D and E). In the anti-*oma-1* piRNAi worms, HRDE-1 ChIP-seq signal at *oma-1* was strongly elevated compared to the one in the control piRNAi worms (Fig. 1E), demonstrating that HRDE-1 is robustly recruited to an artificially induced, nuclear RNAi target gene.

### HRDE-1 interacts with actively transcribed euchromatin in the germline

We previously defined the native nuclear RNAi targets as genomic regions that (1) naturally have high-levels of HRDE-1-bound siRNAs and (2) require HRDE-1 for transcriptional repression and/or heterochromatin formation ^55,56^. These native targets showed no enrichment in the HRDE-1 ChIP-seq signal (*c38d9.2* in Fig. 2A and additional examples in Fig. S1). Although this was contrary to our initial expectation, it can be explained by the finding that the native targets produce very little nascent RNA in wild-type animals ^17^, limiting the recruitment of HRDE-1 to these heterochromatic regions to a level below the detection limit of our assay.

**Figure 2.**
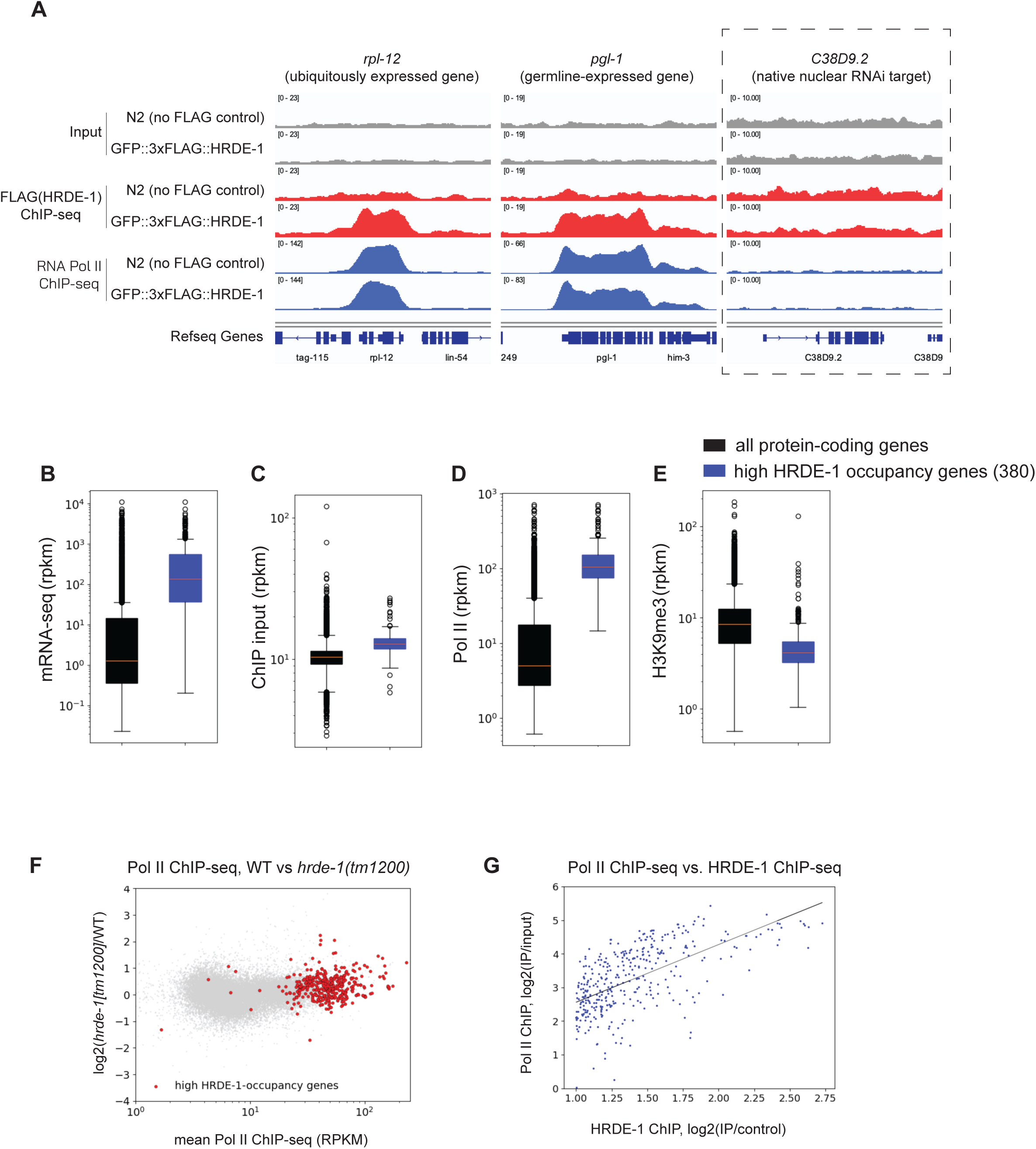
Genes with high HRDE-1-occupancy are actively transcribed. (A) HRDE-1 and Pol II ChIP-seq coverage plots at exemplary high HRDE-1-occupancy genes (*rpl-12* and *pgl-1*) and a control gene *C38D9.2* (a native nuclear RNAi target). (B-E) RNA and ChIP-seq box plots of high HRDE-1 occupancy genes compared with all protein-coding genes. The 380 high HRDE-1 occupancy genes were identified by high HRDE-1 ChIP-seq signals (a minimal of 2-fold enrichment over the control, FDR<=0.05). (F) MA plot comparing WT and *hrde-1(tm1200)* mutant for Pol II levels of all protein-coding genes with high HRDE-1-occupancy genes highlighted. (G) Log2(IP/input) values of Pol II ChIP were plotted as a function of HRDE-1 ChIP for the 380 high HRDE-1-occupancy genes with the least-squares regression line fitted to the data (Pearson correlation coefficient: 0.58, p-value: 6.8×10^−35^).

To identify loci with robust HRDE-1-chromatin association, we performed ChIP-seq peak calling analysis using the MACS3 program ^57^. By using a q-value cuff-off of 0.05 and the broadPeak mode, we identified 1111 genomic regions (Table S1) as HRDE-1-binding regions, with an average size of 2.0 kb. The total HRDE-1-binding regions (2.2 Mb) represent approximately 2.2% of the *C. elegans* genome. The Pol II ChIP-seq signals at these regions were high as well and showed a strong similarity to the HRDE-1 ChIP-seq profiles (Fig. 2A and S2A), We ranked the HRDE-binding regions by the fold enrichment over ChIP input. Regions with highest degree of HRDE-1-enrichment include *csr-1* (5.5-fold enrichment), *prg-1* (5.3), *eft-3* (4.9), and *glh-1* (4.7) (Fig. S2A), which are all known germline-expressed genes ^58^ ^59–61^. Some intronless histone genes such as *his-61* and *his-60* are also strongly enriched for HRDE-1-binding (Fig. S2A). This result indicates that the most prominent HRDE-1-chromatin interaction occurs at transcriptionally active genes.

To further characterize genes associated with HRDE-1-binding, we defined 380 high HRDE-1-occupancy genes as those showing at least two-fold enrichment of HRDE-1 ChIP over the control with FDR ≤ 0.05 (Table S2). These genes exhibit high mRNA abundance, strong RNA polymerase II binding, and low levels of H3K9me3, indicating that they are actively transcribed euchromatic loci (Fig. 2B-E). Approximately 90% of the high HRDE-1-occupancy genes are expressed in the germline, including 18 histone genes, 64 ribosomal protein genes, and well-characterized germline regulators such as *prg-1, csr-1, gld-1, glh-1,* and *pgl-1*.

We next asked whether HRDE-1 binding influences transcription at these highly occupied genes. Re-analysis of published Pol II ChIP-seq and RNA-seq data from synchronized adult wild-type and *hrde-1*(tm1200) mutants ^32^ revealed no substantial changes in Pol II density or mRNA levels for high HRDE-1-occupancy genes, indicating that HRDE-1 has no detectable effect on their expression in adult germ cells (Fig. 2F).

Genome-wide analysis of the WT ChIP-seq samples indicates that HRDE-1 occupancy closely tracks transcriptional activity globally: across the 380 genes, HRDE-1 and Pol II ChIP-seq RPKM (reads per kb per million) values were strongly correlated (Fig. 2G). Finally, HRDE-1 binding is largely restricted to RNA polymerase II transcribed regions, as we detected no enrichment at the RNA polymerase I transcribed 45S pre-rRNA locus or the RNA polymerase III transcribed 5S rRNA gene (Fig. S2B and data not shown). Together, these data indicate that HRDE-1 robustly associates with actively transcribed Pol II chromatin in the germline without measurably altering its transcription.

### HRDE-1-euchromatin interactions are poorly correlated with siRNAs

HRDE-1-bound siRNAs are synthesized by RNA-dependent RNA polymerase (RdRP) EGO-1 and RRF-1 using mRNAs as templates. These siRNAs are usually 22-nt long and have a guanine at the 5’ end, therefore often referred to as 22G-RNAs ^62^. Both nuclear RNAi silenced loci and actively transcribed genes in the *C. elegans* germline generate 22G-RNAs, although they are preferentially loaded into different germline AGO proteins: HRDE-1 and CSR-1, respectively ^56,58^. Nevertheless, a small fraction of the 22G-RNAs from actively transcribed genes are loaded into HRDE-1 ^26,56,63^, raising the possibility that the HRDE-1-euchromatin interaction is guided by these 22G-RNAs. A direct test using siRNA-loading-defective HRDE-1 is not feasible because such mutants fail to accumulate in nuclei ^63^, precluding an assessment of their chromatin binding.

As an alternative, we reasoned that if siRNAs were the primary determinant of HRDE-1 recruitment, the HRDE-1 ChIP-seq signal at high-occupancy genes should positively correlate with the abundance of HRDE-1-associated siRNAs mapped to these genes. Instead, comparison of the HRDE-1 ChIP-seq signals with HRDE-1 co-IP 22G-RNA levels across the 380 high-occupancy genes revealed no detectable correlation (Fig. 3A and S3). This result argues against the guide siRNAs being the instructive signal for the HRDE-1-euchromatin interaction, suggesting an siRNA-independent mechanism behind the interaction.

**Figure 3.**
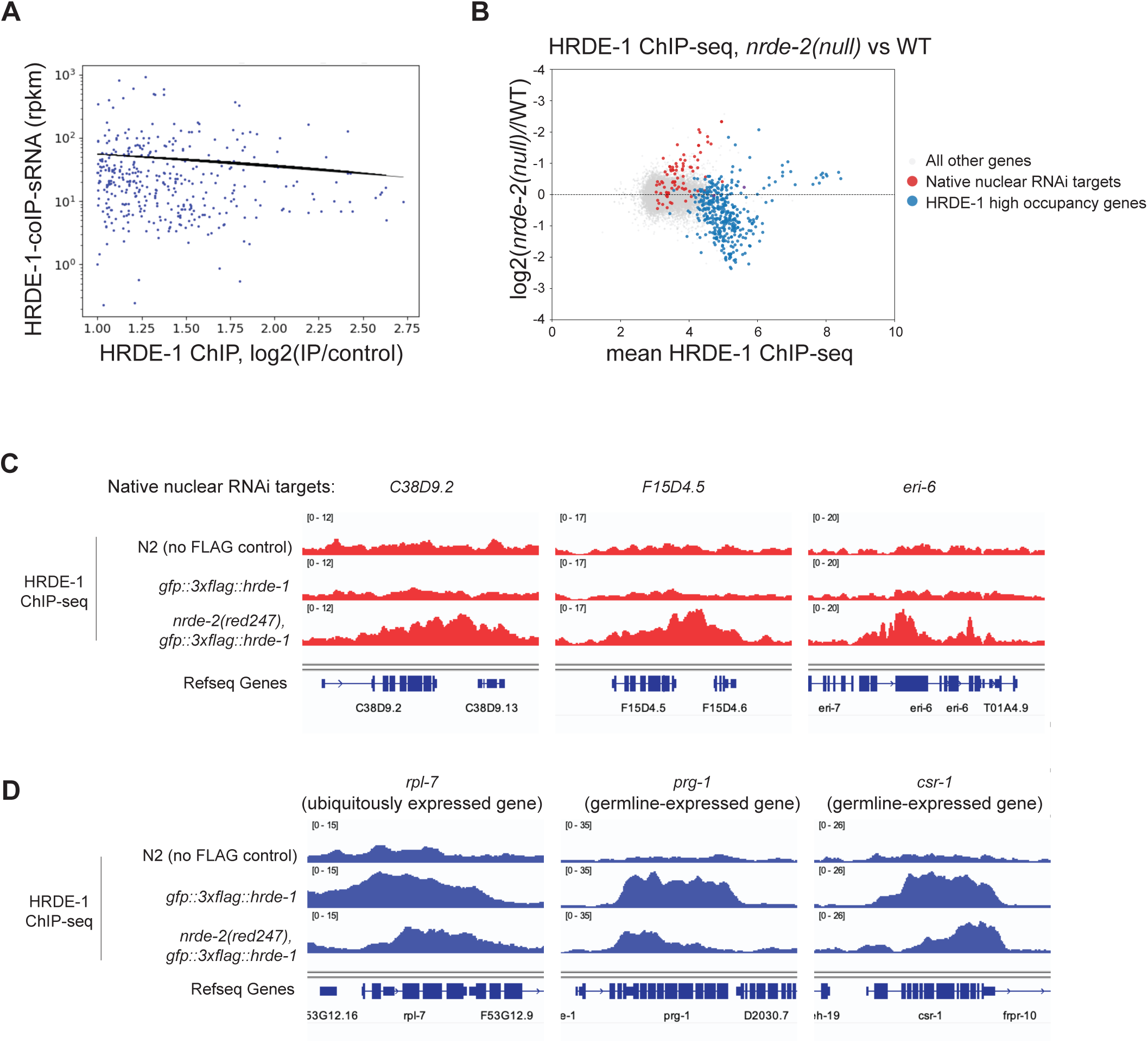
HRDE-1-chromatin interaction does not correlate with HRDE-1-bound siRNAs and can be redirect to native nuclear RNAi targets upon their de-silencing. (A) HRDE-1 coIP siRNAs were plotted as a function of HRDE-1 ChIP for high HRDE-1 occupancy genes with the least-squares regression line fitted to the data (Pearson correlation coefficient: −0.065, p-value: 0.21). (B) MA plot comparing HRDE-1 ChIP-seq between *nrde-2*(WT) vs. *nrde-2(null)* animals. Native nuclear RNAi targets that are transcriptionally silenced by HRDE-1 (red) and HRDE-1 high occupancy genes (blue) are highlighted. (C, D) Coverage plots of HRDE-1 ChIP-seq in *nrde-2* mutant background at exemplary nuclear RNAi targets (C) and HRDE-1 high occupancy genes(D).

### HRDE-1 interacts with chromatin of native nuclear RNAi targets when transcriptionally de-silenced

Despite HRDE-1’s essential role in transcriptional silencing at the native nuclear RNAi targets, its interaction with the chromatin of these targets is relatively low (Fig. S2). This is likely due to the limited production of nascent RNA at these transcriptionally silenced regions. We hypothesize that upon transcriptionally de-silencing of these target, HRDE-1 recruitment to their chromatin would increase accordingly. To this end, we performed HRDE-1 ChIP-seq in *nrde-2* loss-of-function mutant animals. NRDE-2 is a nuclear RNAi factor that functions downstream of HRDE-1, and loss of NRDE-2 results in transcriptional de-silencing of native nuclear RNAi targets ^35,64^. As expected, in the *nrde-2* mutant, HRDE-1 occupancy at nuclear RNAi targets increased significantly (Fig. 3B and C), indicating that HRDE-1 can be efficiently recruited to native targets upon their transcriptional activation. Interestingly, HRDE-1 occupancy at high-occupancy euchromatin genes decreased in the *nrde-2* mutant (Fig. 3B and D). This reduction may reflect redistribution of the available HRDE-1 pool toward the newly transcriptionally active native nuclear RNAi targets.

### The intrinsically disordered RGG domain is required for HRDE-1 chromatin interaction

To pinpoint HRDE-1 regions that mediate euchromatin binding, we first compared its domain architecture and predicted structure to human AGO2. The conserved N, PAZ, MID, and PIWI domains of HRDE-1 closely resemble AGO2, but sequence alignment revealed two HRDE-1-specific insertions spanning residues 1-31 and 304-371 in the N-terminus and PAZ lobe, respectively (Fig. 4A, B and Fig. S4). AlphaFold ^65^ and IUPred ^66^ both predict these segments to be intrinsically disordered regions (IDRs) (Fig. 4B and C). In the predicted 3D structure, the PAZ-proximal IDR projects from the side of the protein opposite to the RNA-binding cleft and forms an apparently flexible surface feature (Fig. 4B). Composition analysis showed that this region is strongly enriched for glycine (28.8%) and arginine (23.5%), with additional enrichment of serine, aspartate, and tyrosine, and contains four RGG and two RG motifs (Fig. 4D). Based on these characteristics, we refer to residues 304-371 as the RGG domain. The RGG domain is well conserved in HRDE-1 paralogs from at least six other *Caenorhabditis* species (Fig. 4E), suggesting a functional importance.

**Figure 4.**
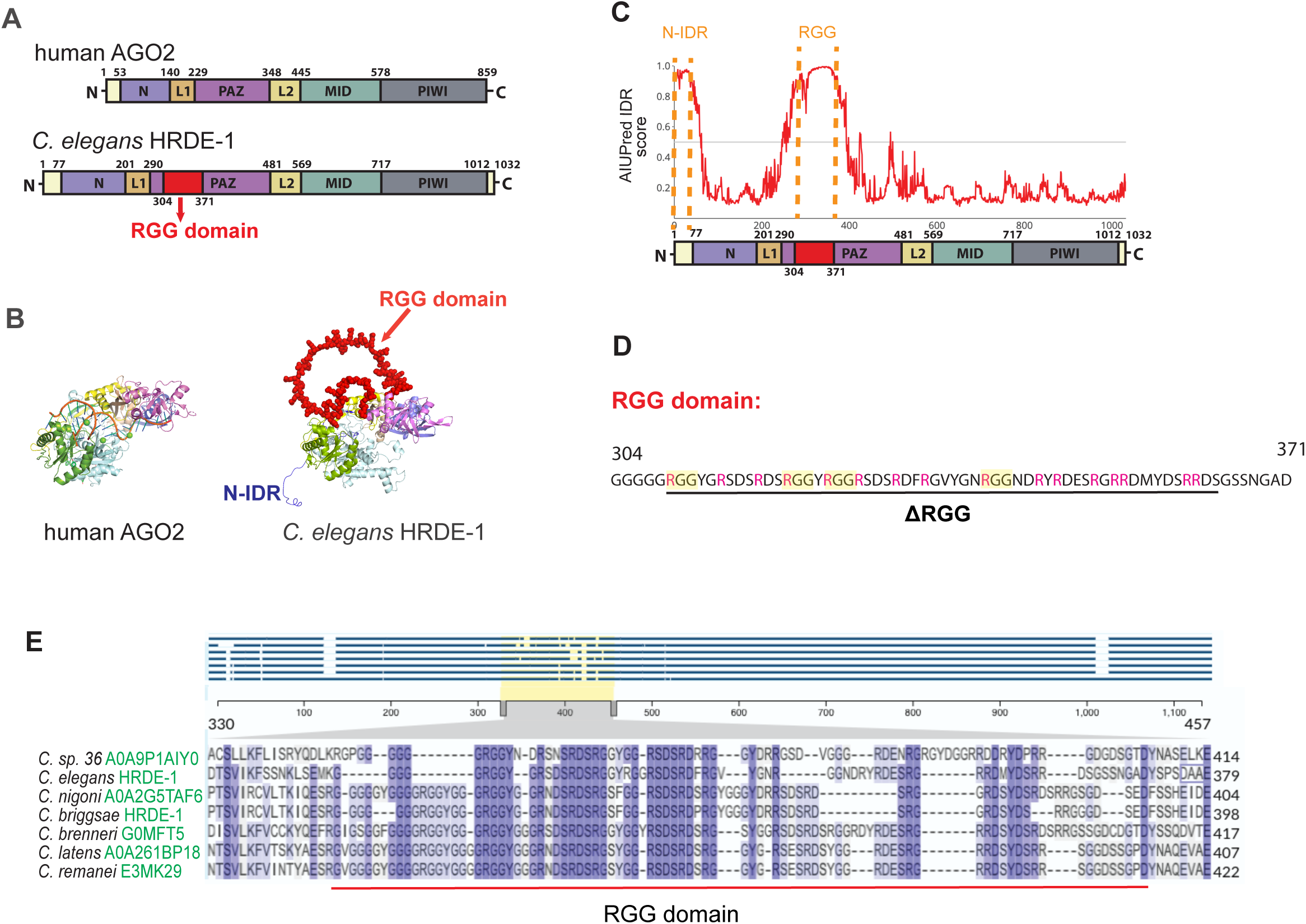
HRDE-1 contains a RGG domain. (A) Domain schematics of human AGO2 and *C. elegans* HRDE-1. (B) 3D structures of human AGO2 with an siRNA::target RNA complex (PDB: 4W5Q ^84^) and *C. elegans* HRDE-1 predicted by AlphaFold ^65^. (C) Intrinsic disorder score analysis of HRDE-1 by AIUPred ^66^. (D) Peptide sequence of the RGG domain from HRDE-1 with the RGG motif and Arg residues highlighted. The peptide sequence deleted in the ΔRGG mutant is indicated. (E) Sequence alignment of RGG domains of orthologous HRDE-1 in different *Caenorhabditis* species (UniProt.org/Align).

To determine the role of HRDE-1’s RGG domain, we deleted the residues 309-364 of the RGG domain in the GFP::3×FLAG::HRDE-1 (named ΔRGG), retaining a short linker to minimize structural distortion (Fig. 4D). We also created a second mutant variant, which deletes the N-terminal IDR (residues 1-51), named ΔN-IDR. The worm strains used in this study carried a germline nuclear RNAi reporter transgene (single-copy insertion) *zc15.3::wrmScarlet*. Both the native *zc15.3* and the *zc15.3::wrmScarlet* transgene are silenced in WT but highly expressed in *hrde-1* loss-of-function mutant ^67^ ^55^. The ΔN-IDR and ΔRGG mutants showed a de-repression of the *zc15.3::wrmScarlet* reporter, indicating a silencing defect (Fig. 5A). Fluorescence imaging revealed that HRDE-1(ΔN-IDR) failed to accumulate in germline nuclei (Fig. 5A), as previously reported ^63^. HRDE-1(ΔRGG), on the other hand, retained the WT-level nuclear localization and robust expression (Fig. 5A). Given our interest in HRDE-1-chromatin interaction, HRDE-1(ΔN-IDR) was not further characterized in this study.

**Figure 5.**
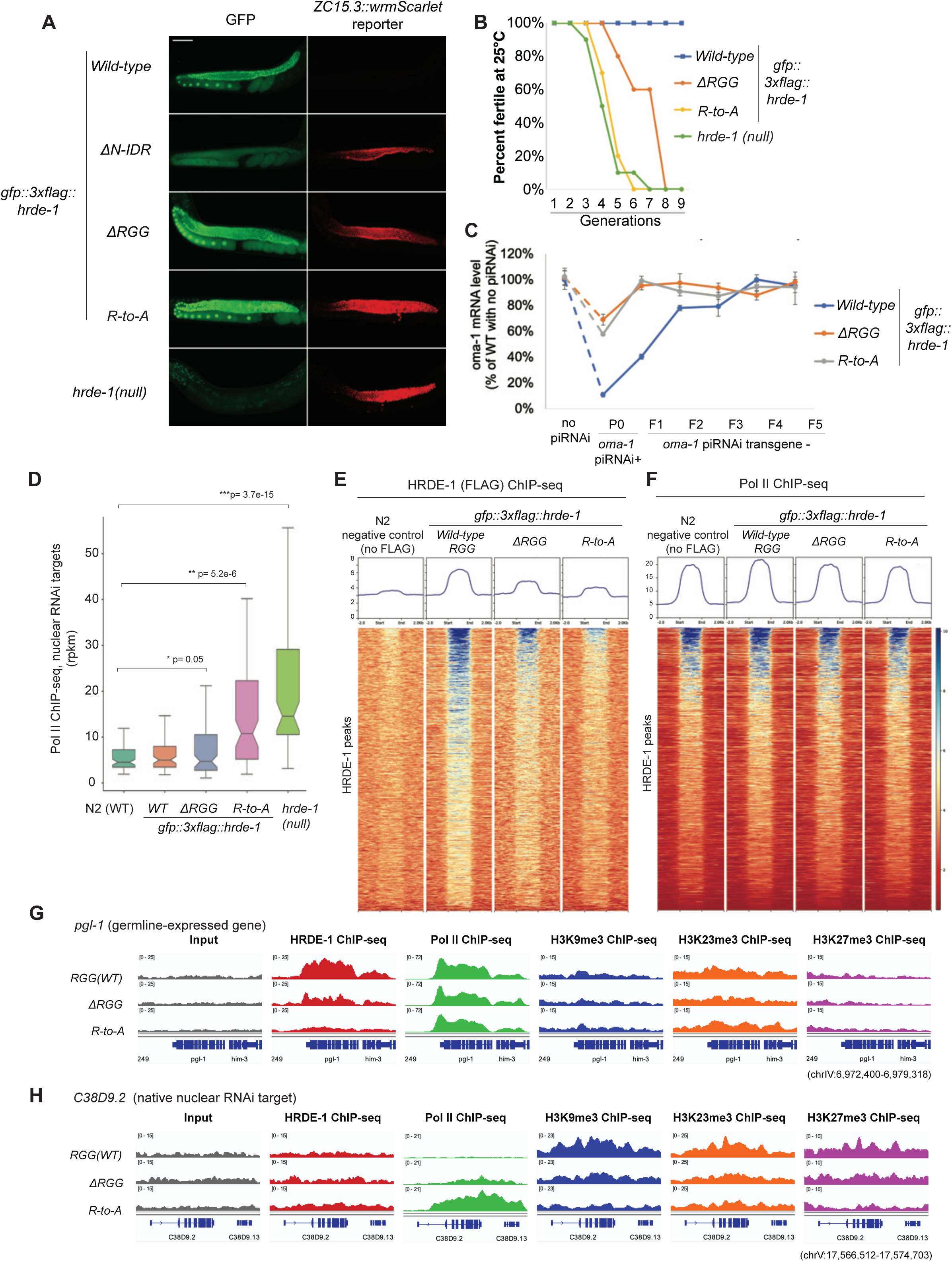
The RGG domain is required for HRDE-1’s functions. (A) Live fluorescent microscopy images of GFP::HRDE-1 and the germline nuclear RNAi report gene (*zc15.3::wrmScarlet*) expressions for in WT and various mutant *hrde-1* adult animals. (B) Transgenerational fertility assay of WT and mutant *hrde-1* animals. Animals were maintained at 16°C before shifting to 25°C for the assay. (C) Heritable piRNAi silencing of *oma-1*. *oma-1* mRNA levels were measured by RT-qPCR. (D) Pol II ChIP-seq box plots of native nuclear RNAi targets (transcriptionally silenced by HRDE-1) in WT and RGG mutants. P values were calculated using paired, two-sided t-test across matched genes. (E and F) Average and pile-up ChIP-seq coverage plots of HRDE-1 (D) and Pol II (E) for 1111 HRDE-1 peaks. (G and H) ChIP-seq coverage plots at *pgl-1* (F, germline gene) and C38D9.2 (G, native nuclear RNAi target) in WT and mutant *hrde-1* animals.

In addition to the silencing defect, we found that the ΔRGG mutant animals displayed a mortal germline phenotype at 25°C, although the ΔRGG mutant population became sterile several generations later than the *hrde-1* null mutants (Fig. 5B). The silencing defect of ΔRGG animals at nuclear RNAi targets was also weaker than the loss-of-function *hrde-1* animals (Fig. 5D), suggesting that RGG deletion does not completely disable HRDE-1’s silencing activity.

HRDE-1(ΔRGG) ChIP-seq analysis revealed a strong defect in chromatin binding at high HRDE-1-occupancy genes compared to wild-type HRDE-1 (Fig. 5E and G). Transcription activity and heterochromatin marks at these genes measured by ChIP-seq remain unaffected by the RGG mutations (Fig. 5F, G, S5B, S5E-F). These results indicate that HRDE-1 interaction with these euchromatin genes require the RGG domain but does not exert an obvious transcriptional silencing effect in adult animals. In contrast, at the native nuclear RNAi targets, RGG mutations resulted in defect in both transcriptional silencing and heterochromatin formation (H3K9/H3K23/H3K27 trimethylation) (Fig. 5D, HS5C-D).

We next asked whether the RGG domain is required for HRDE-1 recruitment and heritable silencing at the exogenous siRNA target. Upon anti-*oma-1* piRNAi, HRDE-1(ΔRGG) failed to accumulate at the *oma-1* locus and the mutant animals exhibited defects in transcriptional repression and H3K9/H3K23/H3K27 trimethylation (Fig. S5A). Transgenerational heritable silencing was also defective in the HRDE-1(ΔRGG) mutant, as measured by RT-qPCR of *oma-1* mRNA in descendants lacking the piRNAi array (Fig. 5C). Thus, the RGG domain is required for HRDE-1 recruitment to both native high HRDE-1-occupancy euchromatin and siRNA-targeted heterochromatin. In addition, the RGG domain is required for the establishment and inheritance of nuclear RNAi (Fig. 5C).

To further probe the contribution of RGG in euchromatin interaction, we created a 2×RGG variant by inserting a second copy of the RGG domain into the WT GFP::3×FLAG::HRDE-1. The 2×RGG variant showed a modest increase in genome-wide HRDE-1 ChIP signal and a WT-level silencing at the reporter gene, further supporting RGG’s role in chromatin binding (Fig. S6). Together, these data indicate that an intrinsically disordered RGG domain in the PAZ lobe is a critical determinant of HRDE-1-chromatin interaction and, consequently, of its function in transgenerational gene silencing.

### Arginine mutations caused more severe phenotypes than the RGG deletion

To test the function of the Arg residues, which have been shown to be essential for RGG-mediated activities in other proteins (reviewed in ^44^), we generated a mutant HRDE-1 in which all 16 arginine residues of the RGG domain were replaced with alanine (R-to-A mutant). We found that, similar to the ΔRGG mutant, the HRDE-1(R-to-A) was able to enter the nucleus but showed defects in gene silencing, transgenerational fertility at 25°C, anti-*oma-1* piRNAi heritable silencing, and chromatin binding (Fig. 5A-E). However, the R-to-A mutant exhibited stronger silencing defects than the ΔRGG mutant, at either anti-*oma-1* piRNAi or the native targets (Fig.5D and S5A). The fertility defect (at 25°C) observed in the R-to-A mutant was also much more severe than the ΔRGG mutant (Fig. 5B). We note that the WT and R-to-A mutant HRDE-1 were expressed at the similar level (Fig. 5A and Fig. S7). These results indicate that the R-to-A mutation caused additional molecular defects compared to the ΔRGG mutant.

### RGG domain is not required for siRNA loading in HRDE-1

To determine whether the RGG domain contributes to siRNA loading, we immunoprecipitated wild-type, ΔRGG, and R-to-A HRDE-1 and analyzed associated small RNAs by 5′ end-labeling and sRNA-seq. End-labeling revealed comparable amounts of co-immunoprecipitated small RNAs for all three proteins (Fig. 6A). sRNA-seq further showed that 22G siRNAs were similarly enriched in wild-type and mutant HRDE-1 complexes and exhibited nearly identical profiles at a native target (Fig. 6B, C). These data indicate that the RGG domain, and its arginine residues, are dispensable for siRNA binding by HRDE-1, consistent with a primary role in chromatin engagement rather than siRNA loading.

**Figure 6.**
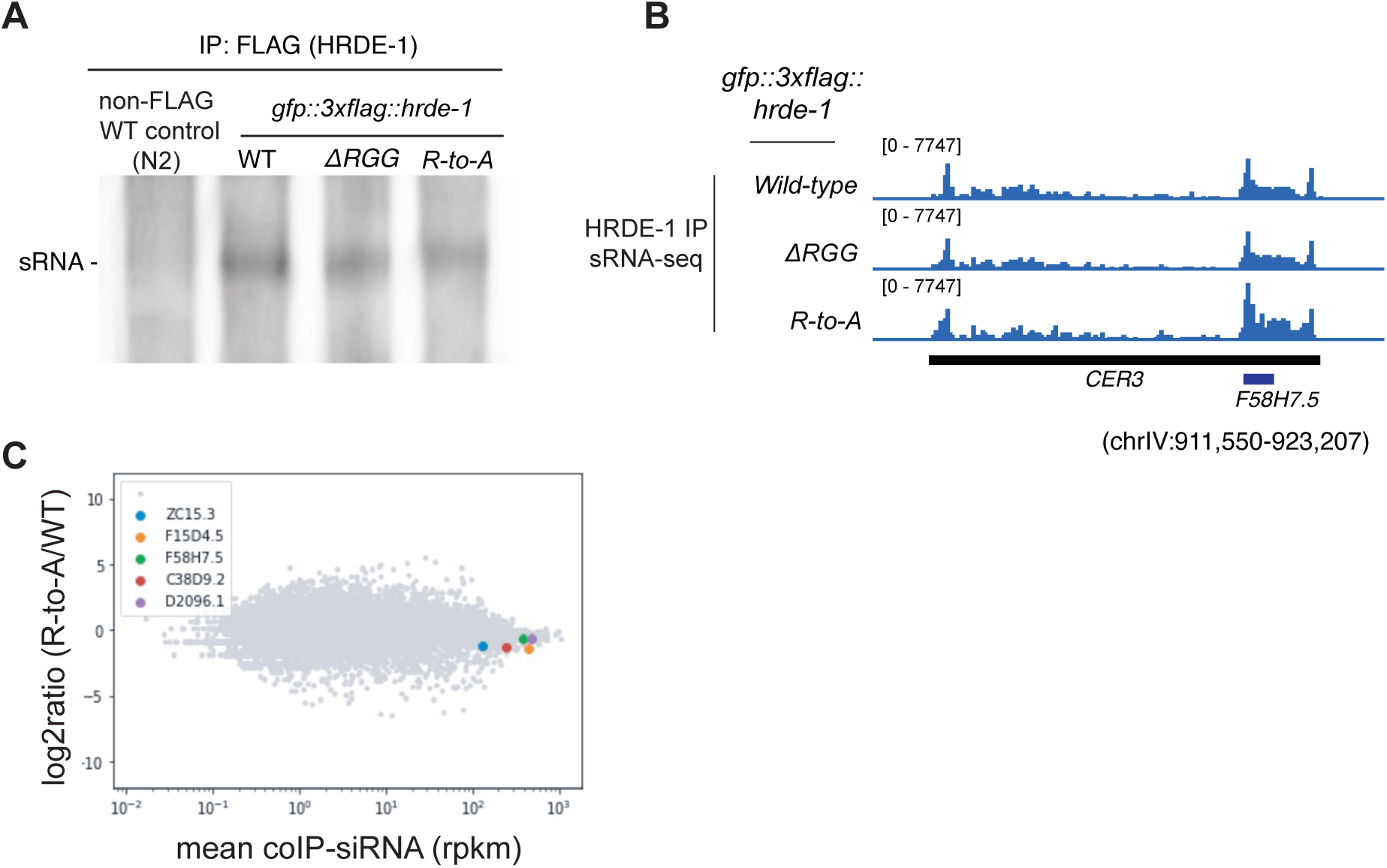
The RGG domain is not required for siRNA binding by HRDE-1. (A) HRDE-1-coIP small RNAs were ligated to biotinylated oligoes, resolved in a UREA-PAGE gel, and visualized by chemiluminescent assay. (B) HRDE-1-coIP small RNA coverage plot at LTR retrotransposon *Cer3*, a native HRDE-1 target. (C) An MA-plot comparing the R-to-A mutant and WT GFP::3×FLAG::HRDE-1 for their HRDE-1-coIP sRNAs of all protein-coding genes. Some of the native germline nuclear RNAi target genes are highlighted with colored dots.

### The function of RGG-domain is dependent on the context

To test whether the RGG domain is sufficient to bind chromatin by itself, we expressed an RGG::GFP::3×FLAG fusion in the germline. A nuclear localization signal sequence was added to the fusion protein, and its nuclear localization was confirmed (Fig. S8A). ChIP-seq analysis showed no enrichment of RGG::GFP::3×FLAG in *C. elegans* genome (an exemplary region shown in Fig. S8B). Thus, the RGG domain by itself is insufficient to confer chromatin binding *in vivo*.

We also created an RGG-relocation mutant by adding the RGG motif to the N-terminus of the GFP::3×FLAG::HRDE-1(ΔRGG). Such RGG-relocation mutant was enriched in the nuclei but remained defective in nuclear RNAi silencing at the *zc15.3::wrmScarlet* reporter, indicating that location of the RGG domain is important (Fig. S8C).

These results together suggest that the RGG function is dependent on additional HRDE-1-specific elements in a context-specific manner.

### RGG is required for the interaction between HRDE-1 and RNA Pol II *in vivo*

Based on the prominent HRDE-1 ChIP-seq signals at the germline euchromatin genes and similar HRDE-1 and RNA Pol II ChIP-seq profiles, we expected that HRDE-1 should interact with RNA Pol II *in vivo* either directly or indirectly. To test this interaction, we performed co-immunoprecipitation(co-IP) experiments. The HRDE-1 and RNA Pol II co-IPed with each other when the animals were treated with the crosslinking agent formaldehyde (Fig. 7), but not when uncrosslinked animals were used (data not shown). Importantly, the HRDE-1-Pol II interaction was defective in both the ΔRGG and R-to-A mutant animals (crosslinked) (Fig. 7). These results indicate that RGG mediates the interaction between HRDE-1 and RNA Pol II, and that the interaction is likely to be weak and transient. We currently do not know whether the two proteins interact directly or through and intermediate factor.

**Figure 7.**
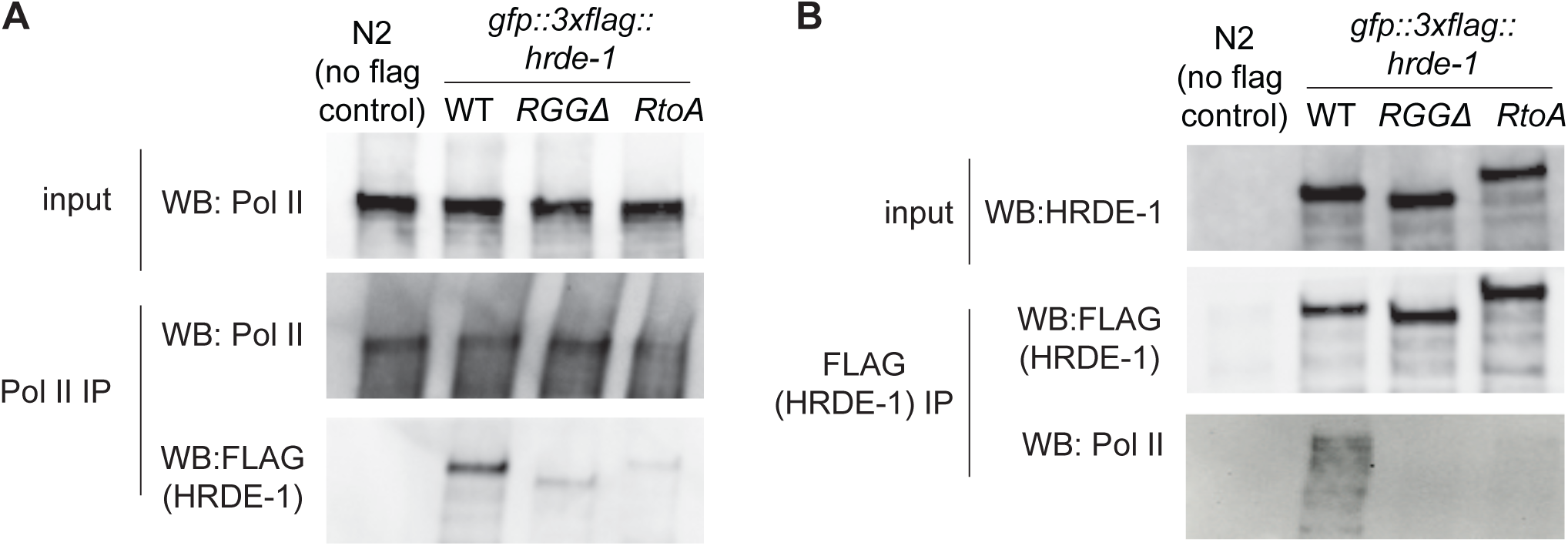
Co-immunoprecipitation analysis of HRDE-1 and Pol II in animals treated with the crosslinking agent formaldehyde. (A) Pol II immunoprecipitation. (B) FLAG (HRDE-1) immunoprecipitation. Pol II and FLAG antibodies used for western blot.

### A heterologous RGG domain from human hnRNP U can functionally replace HRDE-1 RGG

RGG domains are among the most prevalent RNA-binding modules, yet their primary sequences vary widely in length, RG/RGG repeat content, and flanking composition. To test the tolerance of HRDE-1 in RGG sequence change while maintaining a high RGG/RG content, we engineered HRDE-1 chimeras in which the endogenous RGG region was replaced by the following three heterologous RGG domains.

1. The C-terminal RGG domain of human hnRNP U (residues 698-780, Fig. 8A) mediates RNA binding and contributes to chromatin organization in mammalian cells ^46,68,69^. This peptide is enriched for glycine (34.9%), arginine (16.9%), and asparagine (13.3%) and contains seven RGG and five additional RG motifs.
2. We also identified an RG-rich IDR in *C. elegans* somatic nuclear Argonaute NRDE-3 (residues 344-388) that occupies the same structural position as the HRDE-1 RGG domain (Fig. 8A) ^29,30^. It lacks the RGG motif, but is enriched for glycine (22%), arginine (20%), and aspartic acid (13.3%) with an RG and a GR motif.
3. The RGG domain from RNA helicase LAF-1(Fig. 8A), a P granule component in *C. elegans* germ cells, on its own is sufficient to drive liquid-liquid phase separation (LLPS) and form stable condensates *in vitro* and *in vivo* ^70^. This peptide is enriched for glycine (36.2%), arginine (13.4%), and asparagine (13.4%) and contains seven RGG and five additional RG motifs.

**Figure 8.**
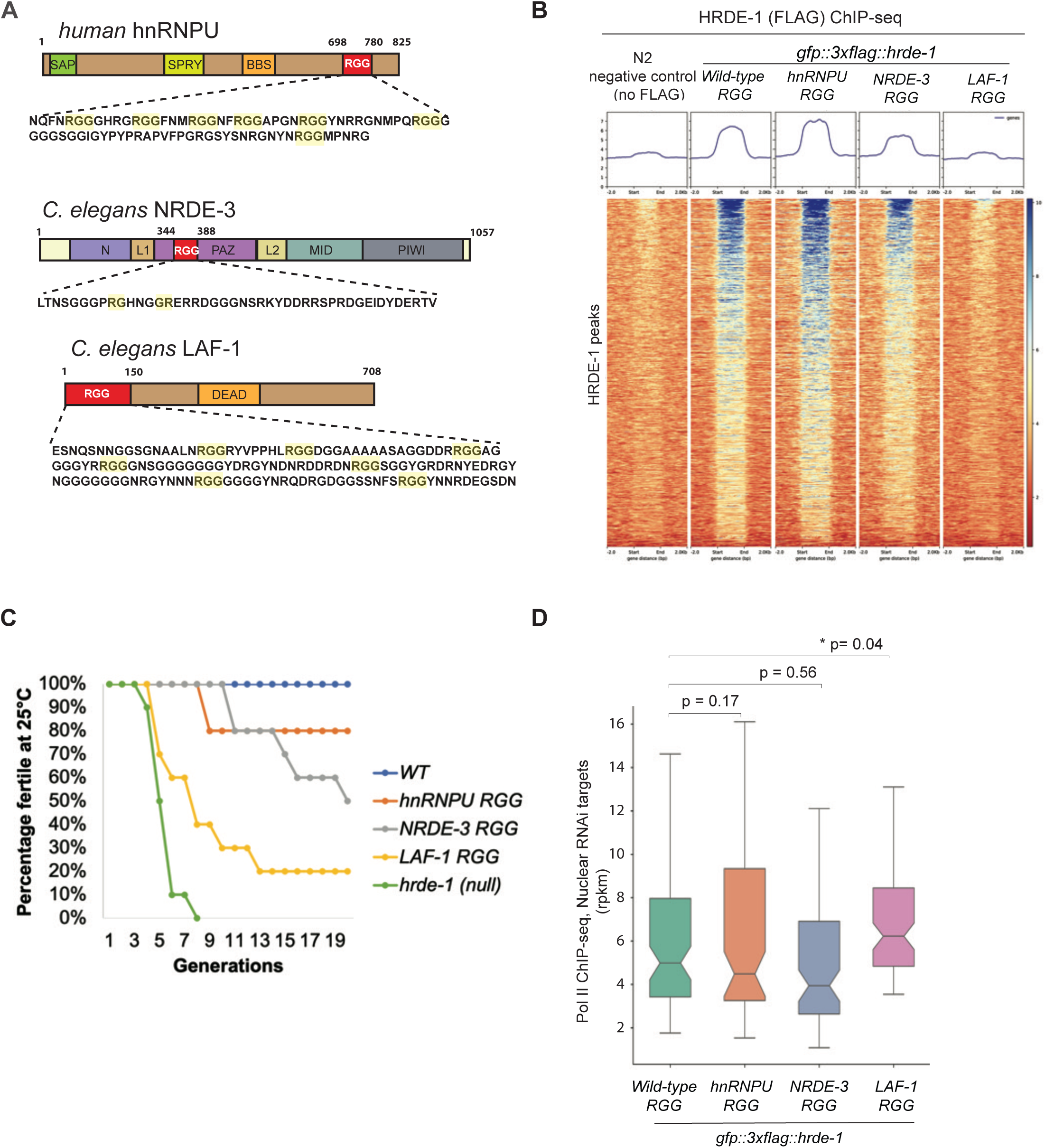
Some heterologous RGG domains can partially rescue the RGG deletion mutant of hrde-1. (A) Schematic diagrams of the RGG and other protein domains in human hnRNP U, *C. elegans* NRDE-3 and LAF-1, with the peptide sequences of the RGG domains shown. (B) Average and pile-up HRDE-1 ChIP-seq coverage plots for the 1111 HRDE-1 peaks. (C) Transgenerational fertility assay of WT and mutant *hrde-1* animals. (D) Pol II ChIP-seq box plots of native nuclear RNAi targets (transcriptionally silenced by HRDE-1) in WT and RGG-replacement mutant animals. P value was calculated using paired, two-sided t-test across matched genes.

All three HRDE-1 chimeras were enriched in germline nuclei (Fig. S9). HRDE-1 ChIP-seq revealed that the HRDE-1-hnRNP U and HRDE-1-NRDE-3 chimera proteins restored near wild-type chromatin-binding profiles at high-occupancy genes (Fig. 8B). Functionally, both chimera strains showed WT-level of fertility at 25°C for 9-10 generations, exhibiting only a very modest mortal-germline phenotype at later generations (Fig. 8C).

The HRDE-1-LAF-1 chimera stain exhibited a stronger mortal germline phenotype than the other two HRDE-1 chimera strains (Fig. 8C). Consistent with these observations, HRDE-1-LAF-1 chimera failed to bind the chromatin of high-occupancy genes (Fig. 8C) and have significant transcriptional silencing defect at the native nuclear RNAi targets (Fig. 8D) compared to the other two HRDE-1 chimeras. These results suggest that the LAF-1 RGG domain lacks the specificity needed for HRDE-1-euchromatin interaction despite its similar RGG/RG content and overall amino acid compositions with HRDE-1.

Given that hnRNP U and NRDE-3 RGG/RG domains have distinct primary amino acid sequences and substantially different number of RGG/RG motifs, yet both can largely substitute for the HRDE-1 RGG domain, these findings suggests that the overall physicochemical properties of the HRDE-1 RGG domain are more critical than its precise primary sequence for promoting HRDE-1-chromatin interaction and transgenerational silencing. RGG-mediated chromatin interaction is likely to be an evolutionary conserved function in a diverse set of molecular pathways.

## Discussion

### The model: a two-step chromatin-targeting mechanism

This study reveals that the nuclear Argonaute HRDE-1 can broadly interact with euchromatin in *C. elegans* germline. We suggest a two-step targeting model for HRDE-1: Step 1, the RGG domain enables HRDE-1 to enter transcriptional active chromatin compartment. Step 2, once inside of the transcribing chromatin compartment, the HRDE-1-associated siRNAs scan the pre-mRNA through base paring interactions for transcriptional silencing. Conceptually, this two-step search strategy is analogous to the mechanisms proposed for Cas9 and transcription machinery, which use transient, low-specificity contacts to scan genomes before establishing stable, sequence-specific interactions ^71–73^.

Many RGG/RG-containing proteins undergo liquid-liquid phase separation and nucleate biomolecular condensates such as P granules, nucleoli, and stress granules (reviewed in ^44^). Chromatin-associated machineries, including heterochromatin complexes, transcriptional hubs, and spliceosomes, also form condensates that organize biochemical reactions in space and time (^41^ ^43^ reviewed in ^73^ ^42^). HRDE-1’s RGG domain may promote its incorporation into, or docking onto, such condensates associated with transcribing genes or pre-mRNA processing, thereby bringing HRDE-1 and its siRNA cargo into proximity with nascent transcripts that would otherwise be occluded within dense RNP assemblies. In this view, RGG-mediated condensate interactions help align HRDE-1 with transcriptional condensates, while the inherently weak and dynamic nature of these contacts explains why HRDE-1-riched euchromatic genes are not silenced in the absence of an RNAi trigger. Stable, siRNA-directed base pairing to nascent transcripts would then be required to transition from a permissive, scanning state to a downstream silencing state that leads to a durable heterochromatin formation and transgenerational repression.

Mutant HRDE-1 that lacks the RGG domain shows a strong defect in silencing, but the phenotype is not as strong as the hrde-1 *null* mutant. We suggest that, in the absence of RGG, HRDE-1-associated siRNAs can still bind its target, but the targeting rate is much lower than the wild-type HRDE-1 due to the barrier activity of the euchromatin condensates.

### Other potential functions of HRDE-1-euchromatin interactions

HRDE-1-euchromatin interactions may also contribute to large-scale chromatin architecture. HRDE-1 has been implicated in chromosome compaction and heterochromatin formation across broad genomic domains ^74–76^. The RG motifs in the N-terminus of CSR-1A has been shown to promote H3K9me3 in *C. elegans* spermatocytes ^77^. Although we observed minimal effects of *hrde-1* loss on transcription of high-occupancy genes in adult germ cells, we cannot rule out that under different developmental stages or environmental conditions, euchromatin-enriched HRDE-1 regulates chromatin organization and gene expression.

The RGG domain itself may provide a regulatory layer through post-translational modification. Arginine residues within RGG/RG motifs can be methylated by arginine methyltransferases, which can tune RNA binding, phase separation, and protein-protein interactions in other RNA-binding proteins (reviewed in ^44^). Future experiments are needed to determine whether the RGG of HRDE-1 is methylated or not. RGG methylation was also shown to promote the stability of PIWI proteins in *Drosophila* ^78^, although neither RGG deletion nor arginine to alanine mutation of HRDE-1 resulted in a detectable protein stability defect. Given the proximity of the HRDE-1 RGG to the small-RNA-binding cleft and its role in RNA- and chromatin association, arginine methylation can potentially modulate HRDE-1’s affinity for chromatin-associated RNAs or downstream protein factors. The R-to-A RGG mutant exhibited more severe defects than the ΔRGG mutant. This is likely caused by some kind of inhibitory effect associated with the mutant RGG domain, supporting a regulatory role of RGG on HRDE-1’s activity.

### Euchromatin interaction as a conserved function of Argonaute proteins

Although Argonaute proteins are best known for their roles in post-transcriptional gene silencing, multiple Argonaute family members in plants, flies, and mammals associate with euchromatin and participate directly in transcriptional or co-transcriptional regulation (reviewed in ^79^). *Arabidopsis* AGO1, which has a RGG domain at the N-terminus, binds the chromatin of transcribed genes and can promote transcription of its targets ^80^. *Drosophila* AGO2 are enriched in the euchromatic bands on polytene chromosomes and has been linked to insulator function and higher-order chromatin topology ^20–22^. In mammalian cells, human AGO1 is enriched at promoters of actively transcribed genes ^23^, and both AGO1 and AGO2 have been implicated in regulating alternative splicing ^24,25^. Mouse AGO2 interacts with open chromatin in male meiotic germ cells and promotes expression of spermatogenic genes ^81^. Together, these studies reveal that chromatin-embedded functions of Argonautes are widespread and extend beyond their well-established post-transcriptional silencing roles in the cytoplasm. It remains unclear whether the euchromatin recruitment of these AGOs mentioned above is mediated primarily by sRNA, additional protein factors, or intrinsic disordered domains. This study provided the first evidence of the involvement of the RGG domains in promoting nuclear AGO chromatin recruitment.

### RGG in Argonaute proteins

We performed a limited survey and found that RGG-containing AGOs are present in human, *Drosophila*, *Arabidopsis*, and *C. elegans* (Fig. S10). Although the RGG domain can be found in the N-terminal, internal, and C-terminal region of many non-AGO proteins ^44^, the RGG domain of AGO proteins found in our survey is limited to the PAZ domain (HRDE-1, WAGO-10, and NRDE-3, all from *C. elegans*) and N-terminus (the rest of the RGG-containing AGOs in Fig. S10). The N-terminus of AGOs is also a frequent location for intrinsic disordered regions other than RGG domains (non-RGG IDRs). Our studies suggested that the location of RGG in HRDE-1 is important to its function. Future studies are needed to determine the mechanism behind the spatial specificity of RGG.

A very limited number of studies, including this one, suggest that the functions of RGG in AGO proteins is diverse. For example, the RGG domain of the *Drosophila* PIWI promotes protein stability through arginine methylation ^78^. The RGG domain of the murine Piwi proteins mediates protein-protein interactions with Tudor proteins ^82^. The RGG domain of the worm CSR-1 protein promotes heterochromatin ^77^. This is consistent with the diverse functions associated with RGGs in non-AGO proteins (reviewed in ^44,45^). RGG may carry out specific functions depending on the location or environment of the hosting protein. It is also possible that the RGG sequences are fined tuned to carry out specific biochemical functions.

Many AGOs do not have any RGG domain or other types of IDRs. For example, the sole AGO in *S. pombe* AGO1, which has been extensively studied for nuclear RNAi, does not have any putative IDR domain. It will be interesting to see if any AGO1 associated protein *in S. pombe* has an RGG domain or other type of IDRs, which may promote the AGO1-euchromatin interaction. Notably, human AGO2 also lacks apparent IDRs, but it interacts with TNRC6B, a member of the GW182 protein family that contains an IDR capable of promoting condensate formation and microRNA-mediated gene silencing ^83^.

### Chimera HRDE-1 with human RGG

As mentioned earlier, the RGG domain is present in a large number of eukaryotic proteins. Although different RGG domains share some common features such as enrichment of certain amino acid residues and peptide motif (RG or RGG), they differ in size, number of RG/RGG motif, amino acid composition, and sequence ^44^. It is very likely that sequences of different RGG domains may have evolved to fit specific functions. In this study, we tested different heterologous RGG domains in their abilities to replace HRDE-1’s RGG domain. Although the scope of this chimera study is limited, several conclusions can be made.

First, RGGs from hnRNP U and NRDE-3 differ from HRDE-1 RGG in the size, RGG/RG motif frequence, and amino acid composition, yet they can partially replace HRDE-1’s RGG, suggesting some degree of degeneracy in RGG’s sequence requirement for euchromatin binding. Notably, HRDE-1 and NRDE-3 are both nuclear AGOs that mediate transcriptional silencing. Though hnRNP U is not a nuclear AGO, it is similarly enriched in the nucleus, and regulate chromatin structure, transcription and co-transcriptional RNA processing in an RGG-dependent manner ^47^. Second, there is still a minor gap between the chimera HRDE-1 with the RGG from hnRNP U or NRDE-3 and the WT HRDE-1 in silencing and fertility, suggesting that specific aspects are optimized for individual RGGs to best fit their functions. Third, RGG from LAF-1 functions poorly in the chimeric HRDE-1 despite a high frequence of RGG motif. LAF-1 is a P-granule component and is not enriched in the nucleus. We suggest that there are different classes of RGG domains that are functionally distinct and incompatible as replacement. In addition, the context of the RGG domain is important, as relocating HRDE-1’s own RGG from PAZ to the N-terminus inactivates HRDE-1. All together, these studies point toward a complex grammar of the RGG domain and highlight the importance of deciphering the RGG code using a tractable system such as HRDE-1 and nuclear RNAi in *C. elegans*.

## Supplemental Figure Legends

**Fig. S1.**
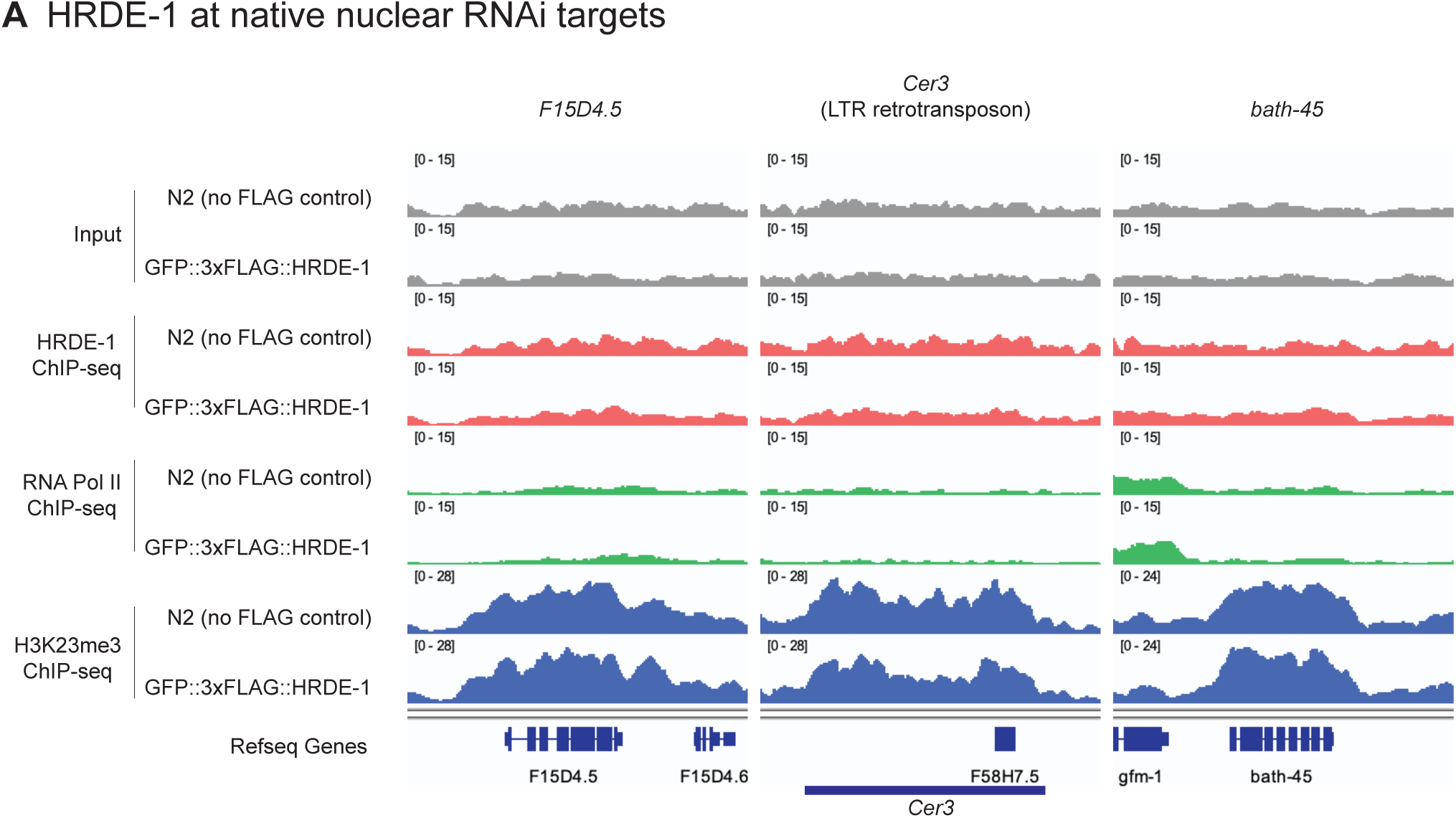
HRDE-1, RNA Pol II, and H3K23me3 ChIP-seq coverage plots at exemplary germline nuclear RNAi targets.

**Fig. S2.**
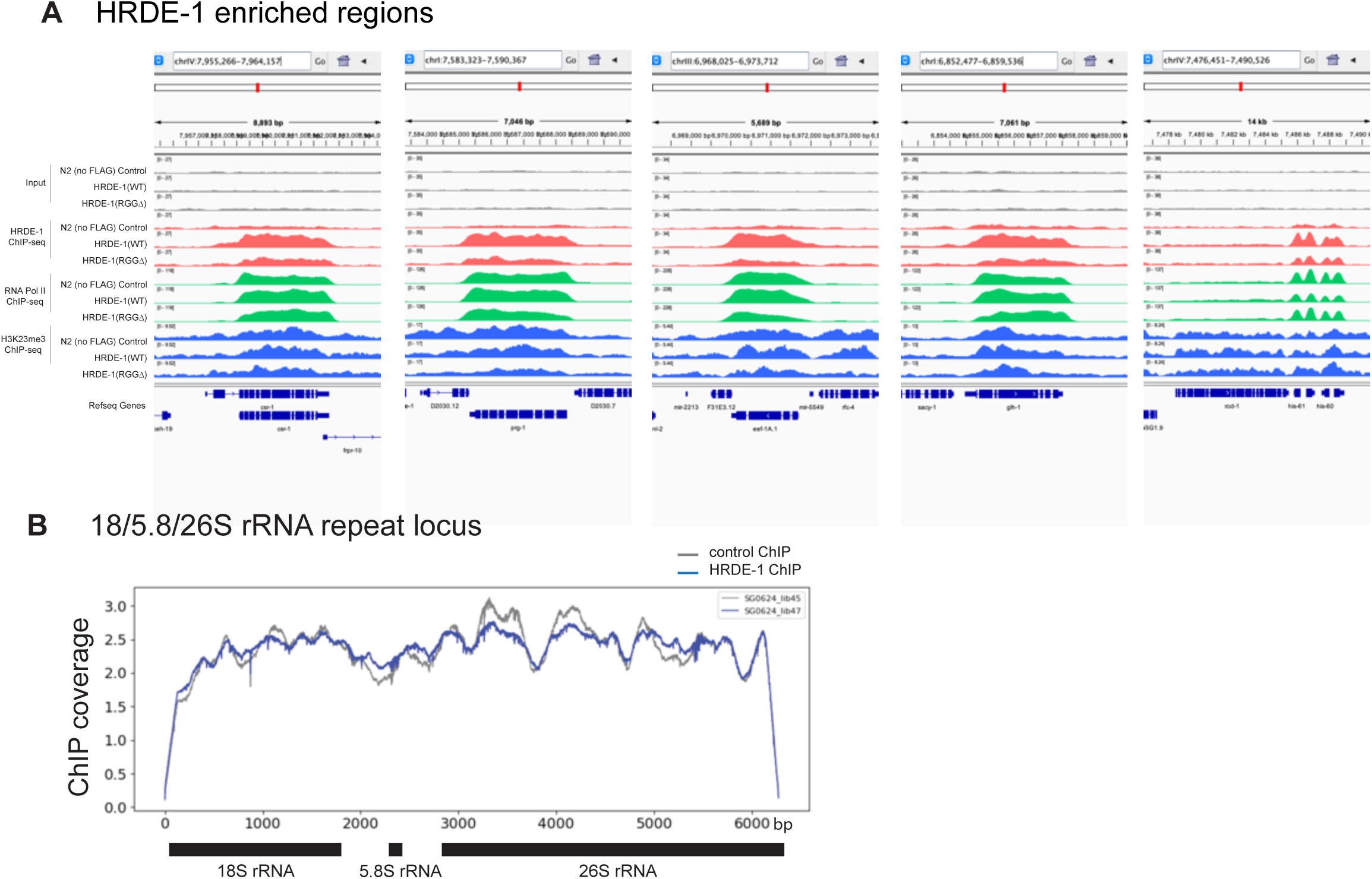
HRDE-1, RNA Pol II, and H3K23me3 ChIP-seq coverages plots at exemplary HRDE-1-enriched genes for N2, WT and ΔRGG GFP::3×FLAG::HRDE-1 animals. (A) and HRDE-1 ChIP-seq coverage at 18/5.8/26S rRNA repeat locus (B).

**Fig. S3.**
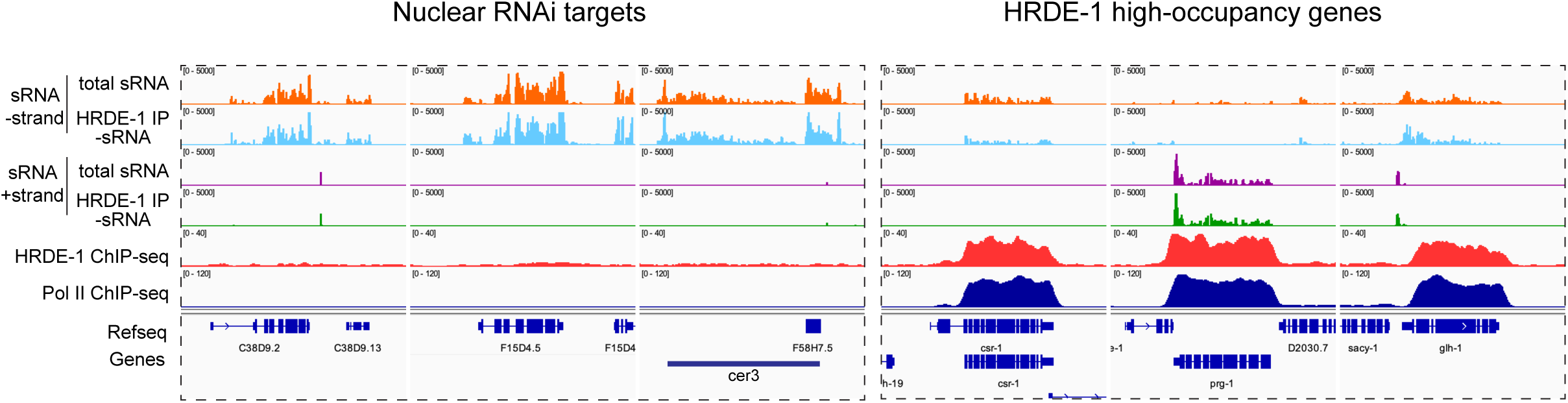
Small RNA-seq, HRDE-1 ChIP-seq, and Pol II ChIP-seq coverage plots at exemplary nuclear RNAi targets and HRDE-1 high-occupancy genes.

**Fig. S4.**
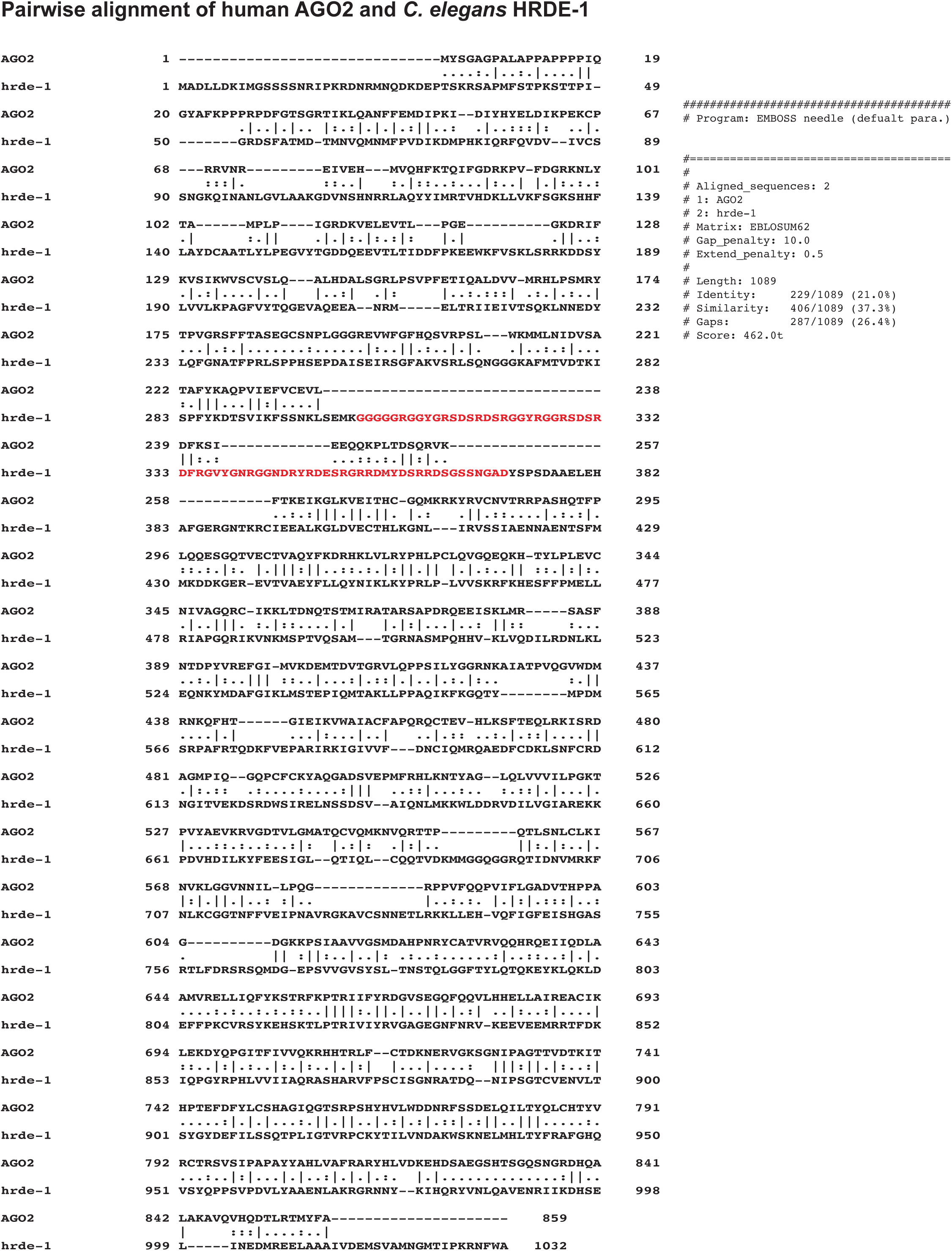
Pairwise alignment of human AGO2 and *C. elegans* HRDE-1 protein sequences. RGG domain is highlighted.

**Fig. S5.**
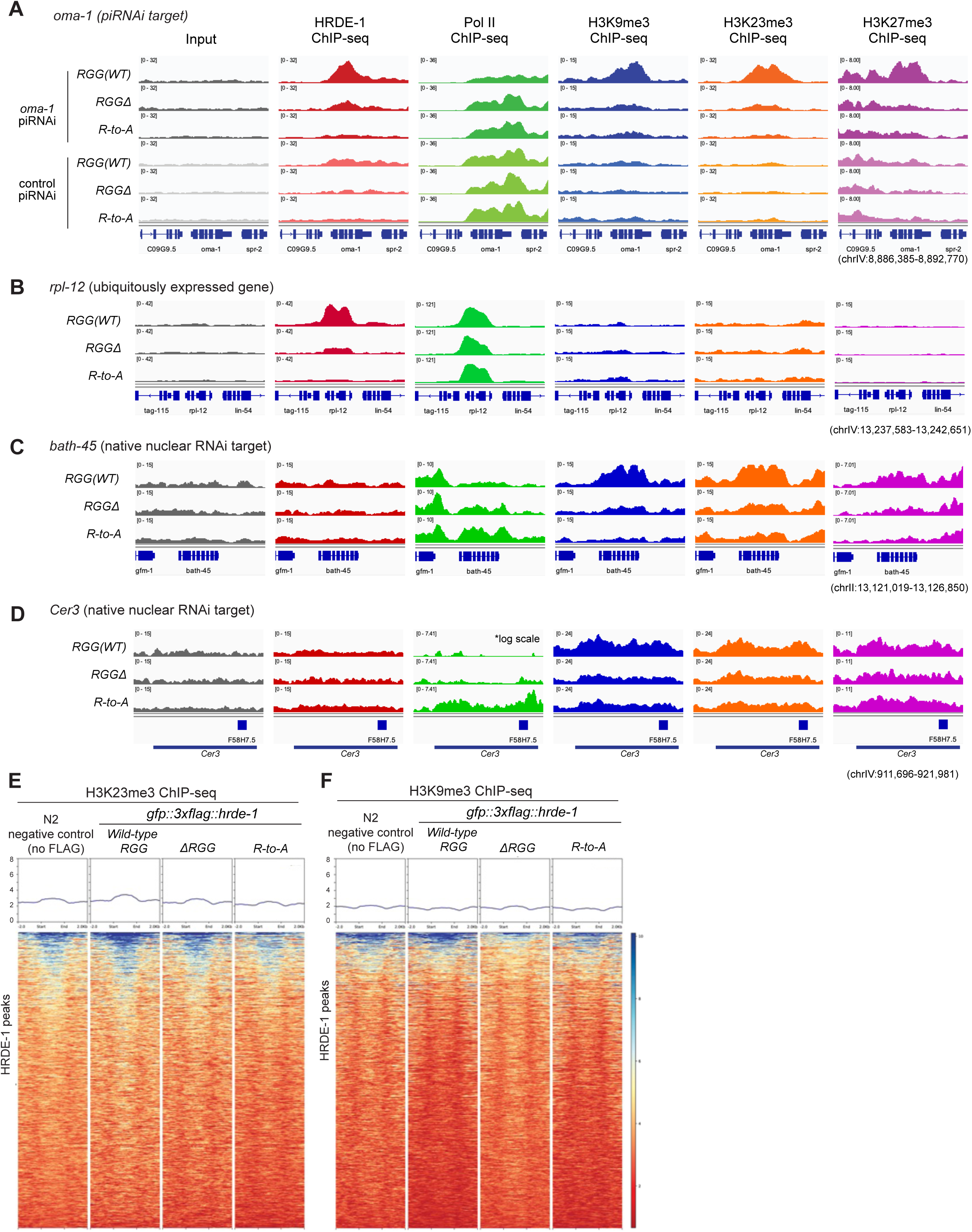
Comparing HRDE-1, Pol II, H3K9me3, H3K23me3, and H3K27me3 ChIP-seq result in WT, ΔRGG, and R-to-A HRDE-1. (A) piRNAi-induced nuclear RNAi is defective in *hrde-1* mutants (ΔRGG and R-to-A). Various ChIP-seq coverage plots were made at the *oma-1* locus in WT, ΔRGG, and R-to-A mutant animals, either with anti-*oma-1* piRNAi or random piRNAi (control). (B-D) Various ChIP-seq coverage plots were made at an exemplary high-HRDE-1-occupancy genes (B) and two native germline nuclear RNAi targets (C, D) in WT, ΔRGG, and R-to-A mutant animals, (E, F) Average and pile-up H3K23me3 (E) and H3K9me3 (F) ChIP-seq coverage plots for HRDE-1 peaks.

**Fig. S6.**
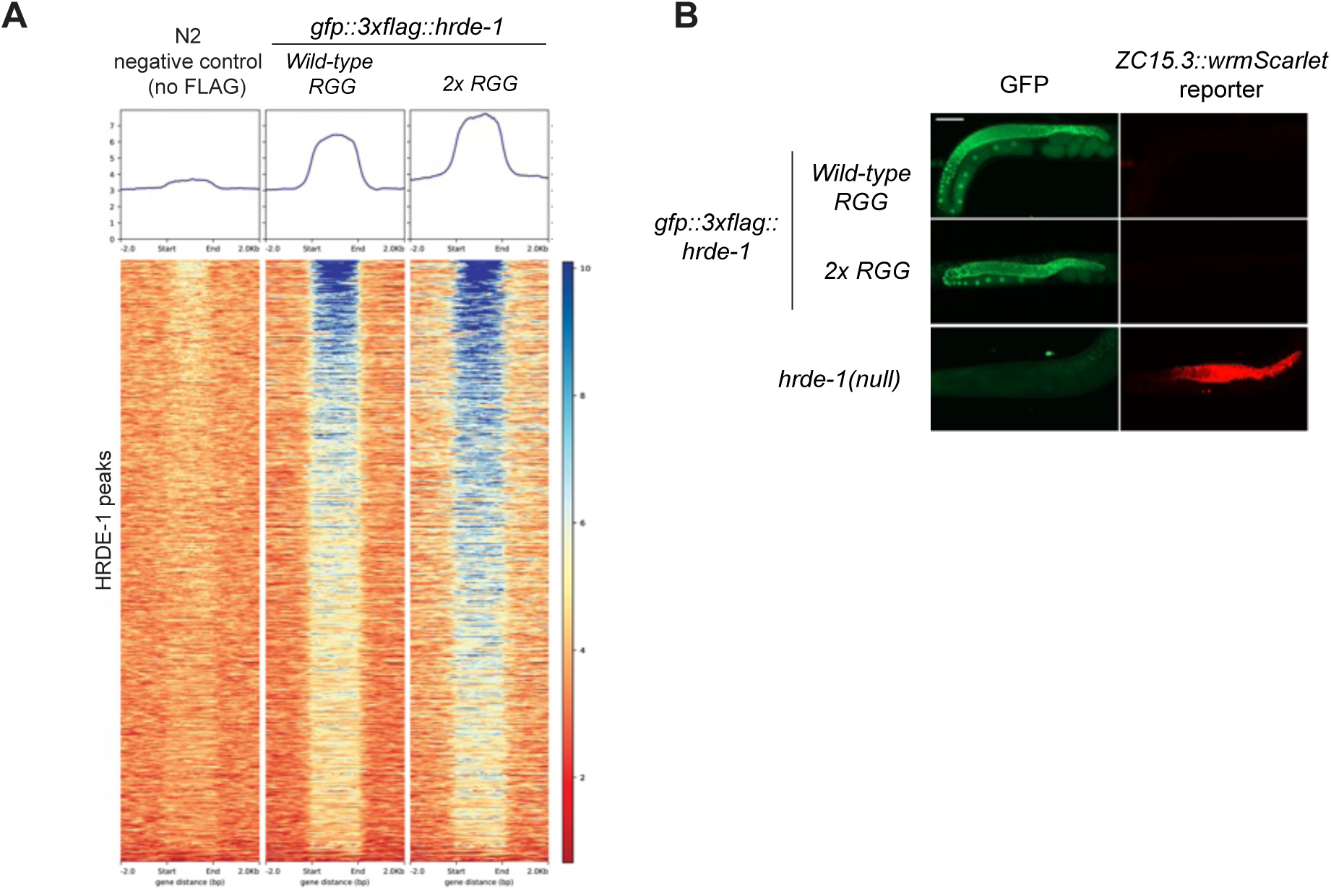
2xRGG enhances HRDE-1-chromatin interaction. (A) Average and pile-up HRDE-1 ChIP-seq coverage plots at HRDE-1 peaks for control, WT, and 2xRGG HRDE-1. (B) Live fluorescent microscopy images of GFP::3×FLAG::HRDE-1 and the germline nuclear RNAi report gene (*zc15.3::wrmScarlet*) expressions for in WT and 2xRGG animals.

**Fig. S7.**
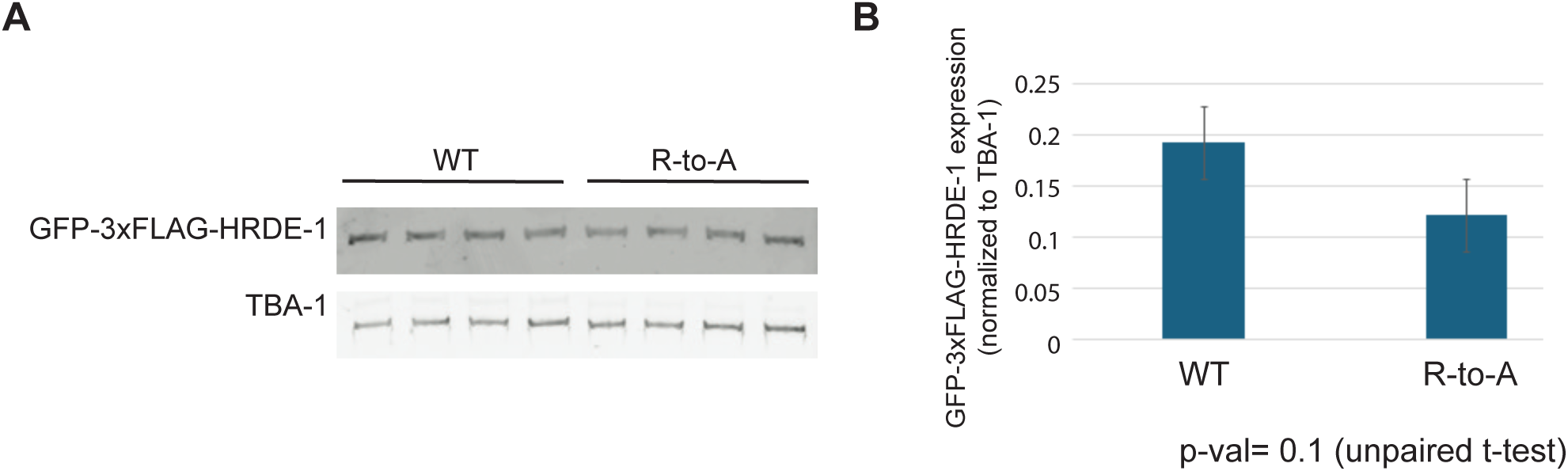
Western blotting analysis comparing WT and R-to-A mutant HRDE-1 expression. (A) Western blot of GFP::3×FLAG::HRDE-1 WT and R-to-A protein by using anti-FLAG antibody (B) Quantification of western blot signal normalized to TBA-1.

**Fig. S8:**
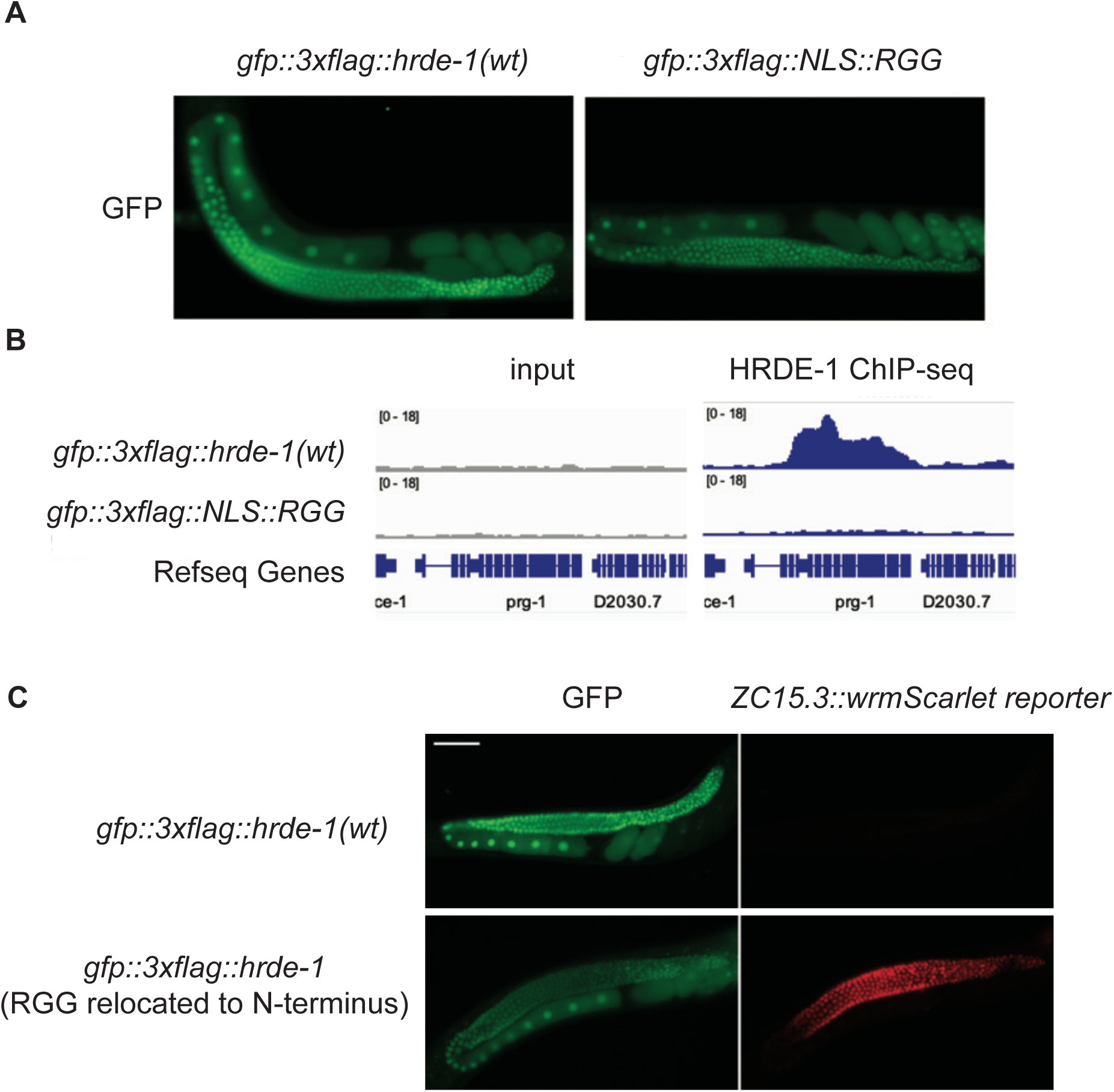
The function of RGG is dependent on its context and location. (A) Live-fluorescent microscopy images of GFP::3×FLAG::HRDE-1 and GFP::3×FLAG::NLS::RGG in adult animals. (B) HRDE-1 ChIP-seq coverage at an exemplary HRDE-1 high occupancy gene, *prg-1*, in the *gfp::3×FLAG::hrde-1(wt)* and *gfp:3×FLAG::NLS::RGG* animals. (C) Live fluorescent microscopy images of WT GFP::3×FLAG::HRDE-1 and mutant protein with RGG relocated to the N-terminus. ZC15.3::wrmScarlet reporter expressions were also shown for the WT and mutant animals.

**Fig. S9.**
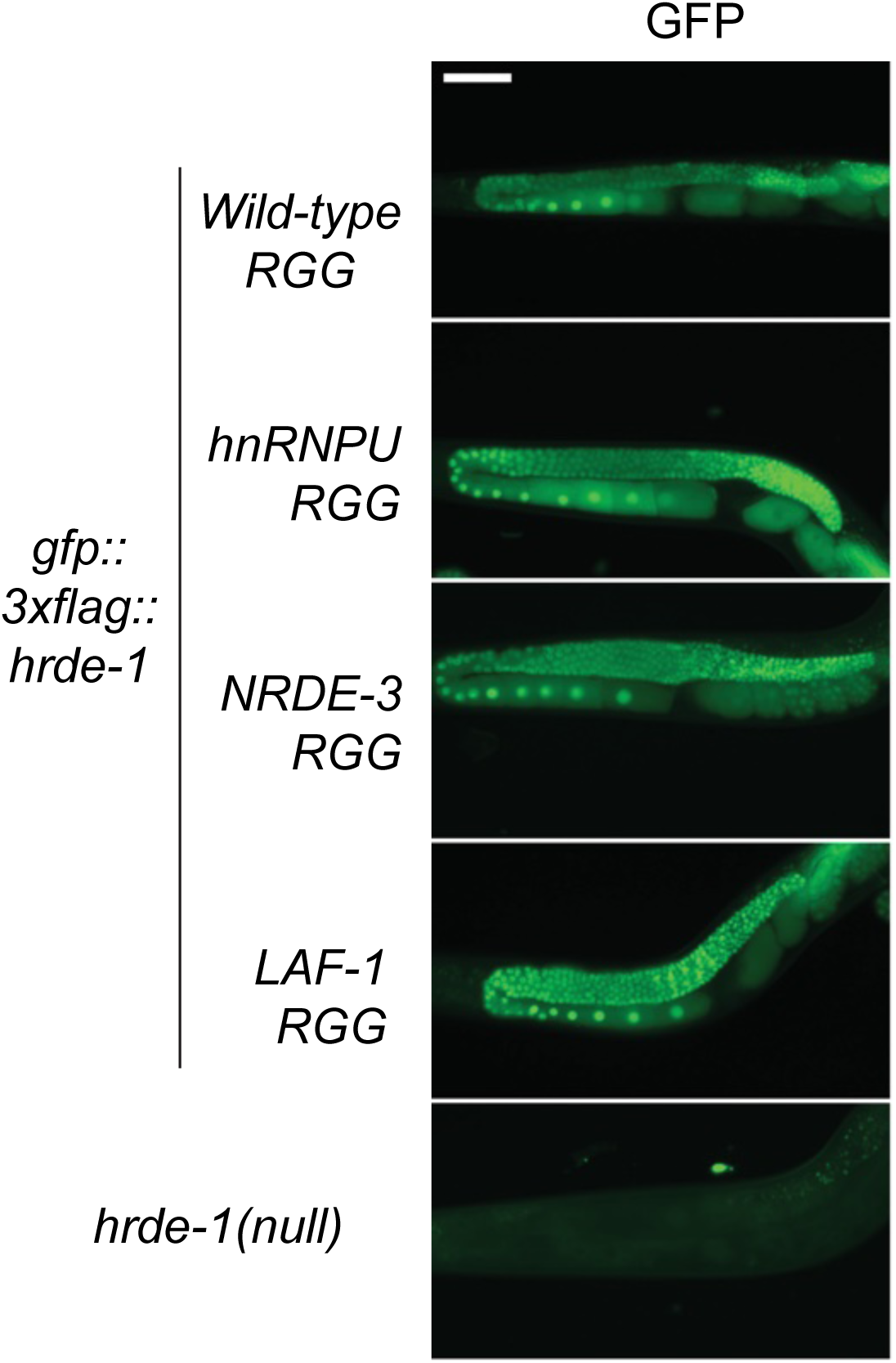
Live fluorescent microscopy images of GFP::3×FLAG::HRDE-1 RGG-replacement adult mutant animals.

**Fig. S10.**
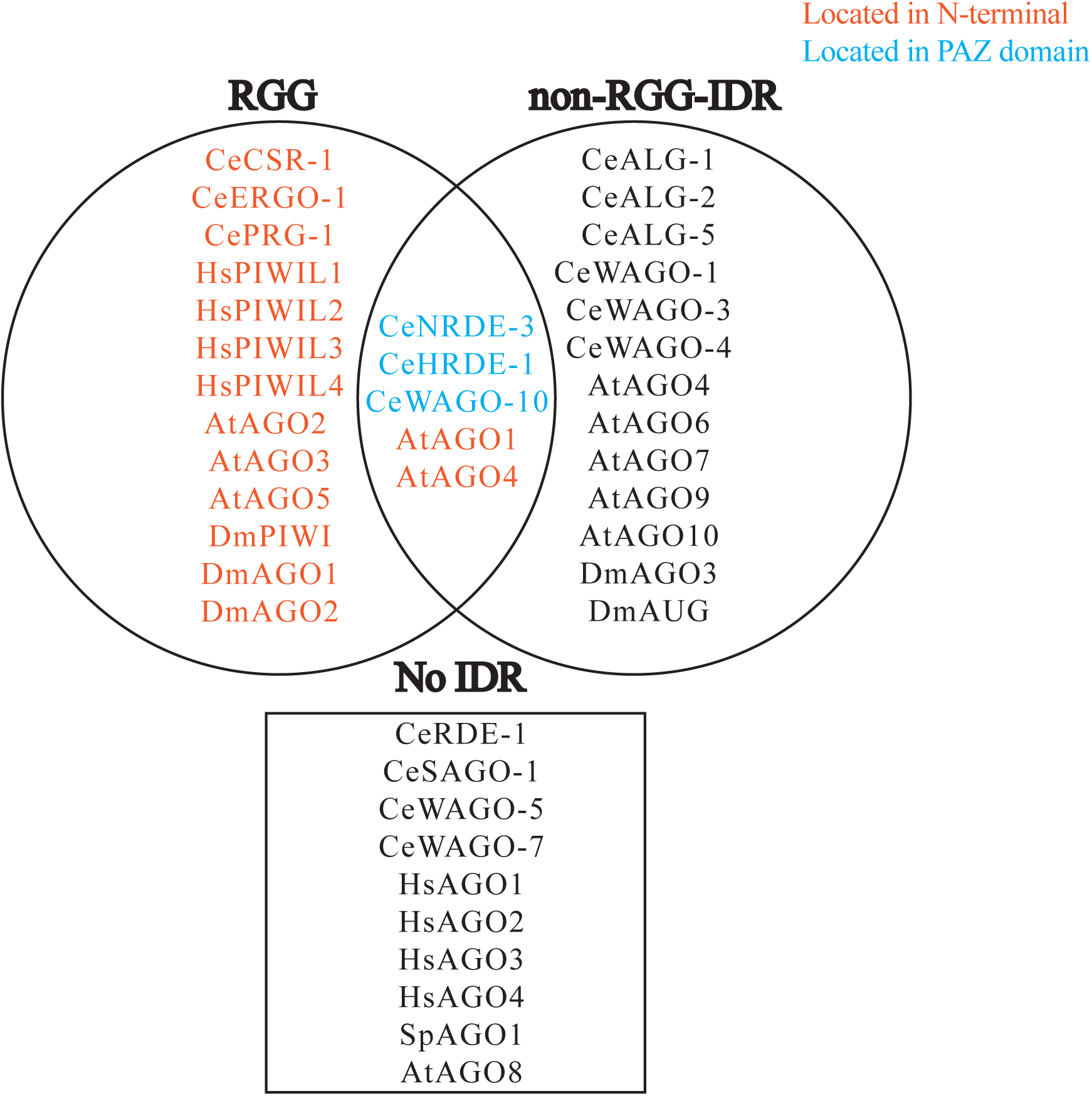
RGG-containing AGOs in human, *Drosophila*, *Arabidopsis*, and *C. elegans*.

## Material and Methods

### C. elegans Strains

*C. elegans* maintenance and genetic crosses were carried out using standard protocols ^85^. Worms were cultured at 20°C on NGM plates with E. coli (OP50). Synchronized young adult animals (68-72 hours after L1 release) were used for experiments unless otherwise indicated. Mutant alleles and strains used in this study were listed in Supplemental Table 3.

### CRISPR-mediated genome editing

HRDE-1 mutations were created by CRISPR-mediated genome editing on the native *hrde-1* gene that carries the *tor125* (GFP::3×FLAG at the N-terminus) allele, created in ^30^. CRISPR crRNA (Synthego) guide sequences and repair template DNA (TWIST Bioscience) sequences were listed in Supplemental Table 3. *dpy-10(cn64)* was used as co-CRISPR maker as described in ^86,87^.

### PiRNAi transgene construction

Strains carrying piRNAi transgenes were generated by DNA injection and extra-chromosomal array formation as described in ^88^. The piRNAi transgene sequences (anti-oma-1 and anti-random DNA as T241 and T119, respectively from ^31^) were listed in Supplemental Table 3. The injection mixture recipe and transgenesis procedure were the same as described in ^32^.

### RT-qPCR analysis

Total RNA was extracted from synchronized adult animals using TRIzol reagent (ThermoFisher) following the manufacturer’s instructions. The RNA was treated with DNase I (NEB) to remove residual genomic DNA and was purified by phenol-chloroform extraction. Complementary DNA (cDNA) was then synthesized using Reverse Transcriptase III (ThermoFisher) according to the manufacturer’s protocol. Quantitative PCR was performed on a QuantStudio 3 real-time PCR system using SYBR Green Master Mix (ThermoFisher). qPCR primers are: oma-1f(SG-1016): 5’-ctacaggactctgcccatacg-3’, oma-1r(SG-1019): 5’-ggcctggcaaacatttctaa-3’, tba-1f(SG-1108): 5’-TCCAAGCGAGACCAGGCTTCAG-3’, and tba-1r(SG-1109): 5’-TCAACACTGCCATCGCCGCC-3’. Relative transcript abundance of *oma-1* was calculated using the ΔΔCT method, with *tba-1* serving as the endogenous control.

### End labeling of HRDE-1 coIP RNA

Adult worms were ground in liquid nitrogen with mortar and pestle and kept frozen in −80°C. Immunoprecipitation (IP) was performed using the FLAG Immunoprecipitation Kit (Sigma, FLAGIPT1-1KT) following the manufacturer’s instructions, with the following modifications. Frozen worm grinds (∼ 400 ul) were resuspended in 600 µl lysis buffer with 6 µl RNaseIn (ThermoFisher Scientific), followed by three times of sonication using Bioruptor (Diagenode) with icy water bath (4°C), 8 minutes each time. The lysate was spin at 13,000 rpm in a bench microcentrifuge for 15 minutes at 4°C and the supernatant was collected for IP and its protein concentration was normalized across samples. 25 µl anti-FLAG M2 gel resin was added to the lysate for 2 hr incubation. After removing the lysate, the gel resin was washed with 500 µl wash buffer three times before the elution step using 0.3 mg/ml 3×FLAG peptide in the wash buffer. IP efficiency was validated by western blotting.

The coIP RNA was extracted using the TRIzol Reagent (ThermoFisher Scientific) and ligated to a biotinylated adaptor using T4 RNA ligase 2, truncated (NEB M0242L). The adaptor sequence was /5rApp/AGATCGGAAGAGCACACGTCTGAACTCCAGTCACAAAAAAAAAAAAAAAAAAAAAAAA/3Bio/(IDT). Ligation products were resolved on an 8% Urea-PAGE gel (National Diagnostics) in 1x TBE running buffer (89 mM Tris-borate, 2 mM EDTA, pH 8.0), then semi-dry transferred to a nylon membrane using the Trans-Blot Turbo system (Bio-Rad) at 0.4 A for 35 minutes in 0.5x TBE. Following transfer, RNA was crosslinked to the membrane using a UV crosslinker (Fisher) at 120 mJ/cm² for 1 minute and detected using the Chemiluminescent Nucleic Acid Detection Module Kit (ThermoFisher Scientific) following the manufacture’s manual.

### *C. elegans* live-imaging

Synchronized young adult worms were paralyzed in 0.1 mM levamisole in M9 buffer (3 g KH₂PO₄, 6 g Na₂HPO₄, 5 g NaCl, 1 ml 1 M MgSO₄, brought to 1 L with H₂O and sterilized by autoclaving), and mounted on 4% agar pads pad. Fluorescence images were acquired using a Zeiss Observer.Z1 epifluorescence microscope equipped with a 20× dry objective and an ORCA-Flash4.0 LT Digital CMOS camera (Hamamatsu). Gonad images were captured with Zen Blue software as z-stacks at 1 μm intervals. The Fiji (ImageJ) software was used to process the images ^89^. To show HRDE-1 and ZC15.3-wrmScarlet cellular localization (Fig. 5), maximum intensity projections were generated from five 1-µm z-stacks of the gonadal side closest to the coverslip. The images in Fig. 5 and Fig. 8 were presented using identical brightness and contrast.

### Multigenerational fertility (mortal germline) assay

Worm strains were kept at 20°C before starting the assay. For the first generation, five L1-stage worms reared at 20°C were moved to fresh plates and incubated at 25°C. Ten plates were set up per strain to provide ten independent replicates. After 6 days, lines with fewer than five progeny on a plate were scored as sterile, whereas lines with at least five progenies were scored as viable and five progenies from these plates were transferred to fresh plates to found the next generation. At every generation, the proportion of viable lines was calculated and used to construct the survival plot.

### Chromatin immunoprecipitation (ChIP)-seq

Chromatin immunoprecipitation (ChIP) and DNA library were constructed as previously described ^55^. Briefly, ∼5,000 young adult worms were grinded in liquid nitrogen and then crosslinked in 2% formaldehyde for 10 minutes at 4 °C. Crosslinking was quenched with 100mM Tris-HCl (pH 7.5). The worm lysates were centrifuged at 8,000 × g for 30 seconds, and the supernatant was sonicated using a Bioruptor (Diagenode) to shear chromatin DNA into 200-500 bp fragments. For each immunoprecipitation (IP) experiment, 2 µg of antibody was added to 400 µl of lysate (containing ∼50 µg DNA) and rotated overnight at 4 °C. Antibodies used in this study are mouse anti-FLAG (Sigma F1804), Rabbit anti-Pol II S2 (Abcam ab5095), Rabbit anti-H3K9me3 (Abcam ab8898), Rabbit anti-H3K23me3 (Active Motif 61499), or Rabbit anti-H3K27me3 (Active Motif 39535). Subsequently, 20 µl of Protein A or G Dynabeads (ThermoFisher Scientific) was added and rotated for an additional 2 hours (Protein A Dynabeads were used for rabbit antibody, Protein G Dydnabeads was used for the mouse antibody). The beads were then washed with high-salt buffer (100 mM Tris-HCl, pH 7.5, 500 mM LiCl, 1% NP-40, and 1% sodium deoxycholate). To elute the immunoprecipitated material and reverse crosslinks, the beads were incubated with 400 µl of worm lysis buffer (0.1 M Tris-HCl, pH 7.5, 100 mM NaCl, 50 mM EDTA, 1% SDS, and 200 µg/ml proteinase K) at 65 °C for 2 hours with agitation every 30 minutes, followed by organic extraction of DNA. The DNA library was prepared by using KAPA HyperPrep Kit (Roche). Each library is uniquely barcoded and were pooled according to the Illumina Novaseq specifications.

### Crosslinking co-immunoprecipitation (co-IP)

Worm grinding, crosslinking, and sonication of chromatin were performed same as the chromatin immunoprecipitation (ChIP) protocol as described in previous section. Antibodies and protein G/A Dynabeads (ThermoFisher Scientific) were added simultaneously to the chromatin lysate and incubated at 4 °C for 2 hours. The beads were then washed three times for 4 minutes each in RIPA buffer (1xPBS, 1% NP40, 0.5% sodium deoxycholate, 0.1% SDS) at 4 °C. To elute the immunoprecipitated proteins and reverse crosslinking, the beads were incubated with 30ul of 2x Laemmli Sample Buffer (BioRad) containing 5% β-mercaptoethanol.at 95°C for 10 mins. Protein samples were separated on a 4-20% Mini-Protean TGX Precast Protein Gel (BioRad) at 200V for 50mins and then transferred to a 0.45 µm nitrocellulose membrane (BioRad) at 25V for 6mins using the Trans-blot Turbo Transfer System (Biorad). Membrane was blocked for 10mins in 3% nonfat dry milk in 1xPBST (1xPBS with 0.1% Tween 20) and then incubated with primary antibody overnight at 4 °C. The primary antibodies used are mouse anti-FLAG (Sigma F1804) at 1:1000, Rabbit anti-Pol II S2 (Abcam ab5095) at 1:1000. Immunodetection was performed using appropriate HRP-conjugated secondary antibodies and SuperSignal West Pico Chemiluminescent kit (ThermoFisher Scienticific), and signals were imaged using a ChemiDoc Imaging System (Biorad).

### RNA-seq library construction

Total RNA was extracted from approximately 1,000 synchronized young adult worms using TRIzol Reagent (Thermo Fisher Scientific), followed by DNase I treatment (NEB) to remove residual genomic DNA. Ribosomal RNA was depleted using RNase H and DNA oligonucleotides antisense to rRNA, as previously described ^17^. The rRNA-depleted RNA was then used to construct barcoded RNA-seq libraries using the SMARTer Stranded RNA-Seq Kit (Takara), following the manufacturer’s protocol.

### Small RNA-seq library construction

For sRNA-seq, total RNA was extracted from ∼1000 synchronized young adult worms using Trizol Reagent (ThermoFisher Scientific) and was treated by DNaseI (NEB). Anti-FLAG immunoprecipitated (HRDE-1 coIP) RNA was directly proceeded to library construction without DNase treatment. The sRNA library construction protocol was adapted from previous study ^38^. Briefly, both total RNA and IP-RNA samples was ligated to 3’ end linker [/5rApp/CTG TAG GCA CCA TCA AT/3ddC/ (IDT)] by T4 RNA ligase 2, truncated (NEB), followed by end processed with Antarctic phosphatase (NEB) and T4 PNK (NEB). The ligation product was size selected using 8% Urea PAGE gel. The purified product was then ligated to a 5’ linker [/5AmMC6/ ACG CTC TTC CGA TCT rXrXrXrX (IDT)], which included a 4nt unique barcode, using T4 RNA ligase 1 (NEB). The ligated RNA was reverse transcribed to cDNA using SuperScript III (ThermoFisher Scientific), and PCR amplified using KAP HiFi kits (Roche).

### High-throughput sequencing and data analysis

Sequencing was performed on Illumina Novaseq X Plus platform using 150nt paired-end reads. All libraries used in this study are listed in Supplemental Table 4. WBcel235/Ce11 was used as the *C. elegans* reference genome for this work. Bioconductor version 3.21 (R version 4.5.1) and Python 3.13.5 were used for the bioinformatic analysis.

ChIP-seq analysis: Rsubread v.1.6.4 ^90^ was used to for the paired-end genomic alignment (parameters: type = “dna”, input_format = “FASTQ”, output_file = bam_out, output_format = “BAM”, nBestLocations= 4, nthreads = 25, phredOffset = 33, unique = FALSE, maxMismatches = 0, indels = 0, PE_orientation= “fr”, minFragLength = 50, maxFragLength = 800, sortReadsByCoordinates = TRUE). A grey list of genomic regions to be excluded from the downstream analysis was generated from the N2 ChIP input by using the greyListBS function (default arguments) in the GreyListChIP package. HRDE-1 ChIP-seq peaks were called by using the MACS3 3.0.3 program ^57^ (“macs3 callpeak -t %s -c %s -f BAMPE --broad -- broad-cutoff 0.05 -n %s --outdir %s --keep-dup auto --max-gap 200 --min-length 200 “, chip_bam, input_bam, output_names, output_names). The overlapping regions between two replicates of HRDE-1 ChIP-seq samples (WT 3×FLAG::GFP::HRDE-1) were considered as HRDE-1 ChIP-seq peaks.

sRNA-seq analysis: sRNA-seq libraries prepared with the 3’-linker ligation method, R1 reads with 5’-barcodes (4 nt) and 3’-linker sequence removed were used for the alignment. The number of perfectly aligned reads (20-24 nt) against the antisense to the mRNA sequences of protein-coding genes were used to calculate the RPKM values for the MA plots.

## Acknowledgement

We thank Zhaorong Zhu, Siddharth Ghosh Roy, Joyce Nadler, James Tang, Vivian Dong, Elaine Gavin for technical assistance. Some strains were provided by the CGC, which is funded by NIH Office of Research Infrastructure Programs (P40 OD010440). Research reported in this publication was supported the National Institute of General Medical Sciences of the National Institutes of Health (R01GM111752 and R35GM152219) to S.G.

